# Attention-Guided Multimodal Neuroimaging Fusion Network for Modeling Brain Aging Pattern

**DOI:** 10.64898/2026.03.28.713645

**Authors:** Zhuo Wan, Wanxiang Fu, Javed Hossain, Leonardo L. Gollo, Kaichao Wu

## Abstract

Brain age prediction from neuroimaging data provides critical insights into neurodevelopmental trajectories and neurodegenerative processes. However, effectively leveraging complementary structural and functional brain information for accurate prediction remains a major challenge. In this study, we propose an Attention-guided Multimodal brain Age prediction Network (AMAge-Net), a novel framework that integrates resting-state functional MRI (fMRI) and structural MRI (sMRI) to enhance brain age estimation. In AMAge-Net, functional features are captured from fMRI through a hierarchical Graph Attention Network, while structural features are learned from sMRI via a 3D DenseNet architecture. To enable effective cross-modal integration, AMAge-Net incorporates a Multi-Head Cross-Attention mechanism followed by a Gated Fusion Module, allowing the model to dynamically prioritize the most informative features from each modality, thereby improving interpretability and predictive accuracy. Evaluation on the Cam-CAN dataset (652 participants, aged 18–89) demonstrates that AMAge-Net outperforms state-of-the-art unimodal and multimodal baselines, achieving a mean absolute error (MAE) of 5.09, root mean square error (RMSE) of 6.52, *R*^2^ of 0.87, and Pearson correlation (PCC) of 0.94. The proposed model further demonstrates robust generalization, achieving an MAE of 4.29, RMSE of 5.59, *R*^2^ of 0.58, and PCC of 0.77 on the independent OASIS-3 dataset. Comparative and ablation studies further confirm the effectiveness of the proposed fusion strategy and modality-specific encoders. Beyond predictive performance, AMAge-Net highlights interpretable brain regions that provide insights into the mechanisms of functional and structural brain aging, while gender-specific analyses reveal distinct aging trajectories between males and females. These findings establish AMAge-Net as a powerful and interpretable approach to brain age estimation, advancing efforts to characterize healthy aging and detect early deviations associated with neurological and psychiatric disorders.

**Author summary:** Estimating the biological age of the brain from imaging data offers a window into normal development, healthy aging, and the early stages of disease. A major challenge is how to combine information from structural scans, which show brain anatomy, and functional scans, which capture brain activity. Here, we present a new computational framework that integrates both types of data to improve the accuracy and interpretability of brain age prediction. Applied to two independent, large-scale lifespan magnetic resonance imaging datasets of individuals spanning early adulthood to late life, our framework produced highly accurate predictions and consistently outperformed existing methods. Beyond predictive performance, the model highlighted brain regions that appear especially important for age-related changes, and it revealed distinct aging patterns between men and women. These findings provide a powerful and interpretable tool for studying how the brain changes across the lifespan, with potential applications in detecting early deviations linked to neurological and psychiatric disorders.

## Introduction

Brain age reflects the biological state of the brain that may diverge from one’s chronological age due to factors such as genetic predisposition, environmental exposures, and lifestyle choices [1, 2]. Early work revealed that cognitive decline and structural brain alterations—such as cortical thinning and white-matter degradation—do not always track with years lived [3–6], implying that the brain follows its own aging trajectory shaped by neurobiological and psychosocial factors. In this regard, brain age prediction driven by neuroimaging has been widely used in studies of brain development and aging [4, 7–11], owing to its high accuracy in assessing brain maturation, aging trajectories, and aiding disease diagnosis [5, 12–14, 14–17]. With the emergence of large-scale neuroimaging and longitudinal cohort studies, researchers have leveraged comprehensive datasets to characterize normative aging trajectories and quantify deviations between biological and chronological age [4, 18, 19]. The difference between predicted brain age and chronological age, known as the Predicted Age Difference, serves as an important indicator for identifying abnormal brain development [8, 10, 20, 21]. A significant difference may suggest accelerated brain aging or delayed maturation. These efforts have established brain age as a powerful integrative biomarker for the early detection of neurodegenerative processes and the prediction of cognitive outcomes in older adults. As global demographics shift toward an aging society, accurately predicting brain age has become critical for guiding timely, personalized interventions—and it is precisely this challenge that our multimodal framework tackles.

Traditional brain age prediction approaches have long depended on regression based and machine learning techniques to extract handcrafted imaging features, reduce feature dimensions, and make predictions with support vector regression [22–24], Gaussian process regression [25, 26], etc. With structural MRI (sMRI) data, these methods extract features like cortical thickness, gray matter volume, and white matter integrity to estimate brain age [5, 27–30], though these methods can suffer from feature-selection bias and limited generalizability [8, 31, 32]. Studies on brain age prediction with functional MRI (fMRI) apply similar methods. Functional connectivity networks (FCNs) were constructed by extracting time series from predefined brain regions or the whole brain. These networks were then used as input features for various machine learning models to make brain age prediction [33–36]. Recently, deep neural networks have become prevalent in manifold neuroimaging studies, including brain age estimation [15, 18, 26, 29, 32, 37–43]. The advent of deep learning has ushered in powerful alternatives: 3D convolutional neural networks autonomously learn multiscale representations directly from volumetric MRI data [29, 38–41, 43–46], while transfer-learning strategies can leverage large, publicly available neuroimaging repositories to then pretrain models that can be fine-tuned on smaller, specialized cohorts. Beyond static anatomy, recurrent neural networks (RNNs) and graph neural networks (GNNs) have been employed to model the temporal dynamics of fMRI time-series and the complex topology of FCNs, respectively [47–52]. More recently, attention mechanisms and transformer-based architectures [53–59] have been introduced to highlight the most informative brain regions and connectivity patterns, improving both interpretability and predictive accuracy.

Despite the progress, these advances have largely unfolded within single modalities—either structural or functional—thereby overlooking the rich, complementary nature of information that is encoded in different neuroimaging modalities when considered jointly [60, 61]. In the context of brain aging, this complementarity arises because structural and functional MRI capture distinct but interrelated aspects of age-related brain changes. Specifically, sMRI provides relatively stable, high-resolution information about long-term anatomical alterations, such as volumetric loss, cortical thinning, and microstructural degeneration, which reflect cumulative effects of aging and neurodegeneration [5, 62]. In contrast, fMRI characterizes more dynamic properties of brain organization, including network coherence, temporal synchrony, and resting-state functional interactions, which may reorganize earlier or more flexibly in response to aging-related processes and can precede overt structural decline [33, 63–66]. As a result, these two modalities encode age-related signals at different spatial, temporal, and biological scales. Previous studies have shown that fMRI- and sMRI-derived features capture partially non-overlapping aging patterns, and that their integration can improve brain age prediction performance [43, 67–69]. By jointly modeling structural constraints and functional reorganization, multimodal fusion has the potential to uncover synergistic biomarkers that are not accessible from either modality alone, thereby refining brain age estimates and increasing sensitivity to subtle pathological deviations

Building on this insight, in this study, we propose an Attention-guided Multimodal brain Age prediction Net (AMAge-Net) framework that integrates sMRI with advanced functional network features derived from fMRI to boost both the accuracy and interpretability of brain age prediction. Specifically, we address three key contributions: First, we introduce a dual-pathway framework for learning that extracts anatomical features from sMRI using a DenseNet121-based [70] network operating on 3D volumetric inputs, with 3D features projected into a compact representation only at the final embedding stage [71], rather than through patch extraction. In parallel, the functional pathway analyses 4D fMRI data (three spatial dimensions plus time) by converting functional connectivity (FC) metrics into graph representations for feature learning. By learning these two representations independently, the dual-pathway framework captures complementary anatomical and functional information that reflects distinct but related mechanisms of brain aging. Second, to overcome limitations in conventional fMRI processing, our framework incorporates a graph attention mechanism that models the FC matrix as a graph, capturing non-local inter-regional relationships and enriching the representation of neural dynamics. Finally, in the multimodal feature fusion section, we employ a Multi-Head Cross-Attention mechanism and Gated Fusion to selectively emphasize discriminative features across modalities, addressing the challenge of noisy or redundant inputs in high-dimensional neuroimaging data and improving age prediction accuracy.

Our results demonstrate that integrating sMRI and fMRI data significantly improves the accuracy of brain age prediction. Beyond that, we employ the saliency map to obtain the structural feature importance and derive the functional feature importance from integrated attention weights. The salient score integrates structural and functional features, highlighting the critical brain regions implicated in accelerated aging, and also reveals the different aging in different genotypes. Together, these benefits underscore the framework’s potential to deepen our understanding of neurodegenerative processes and to enable targeted early interventions for individuals at heightened risk of accelerated brain aging.

## Materials and methods

### Participants and Data Preprocessing

This study used neuroimaging data from the publicly available Cambridge Centre for Ageing Neuroscience (Cam-CAN) dataset [72, 73], which provides a comprehensive resource for studying cognitive aging in healthy individuals. Specifically, this paper used structural T1-weighted (T1w) MRI and fMRI data from 652 healthy participants (322 males, 330 females) with a mean age of 54.9 ± 18.6 years and an age range of 18.5–88.9 years. The mean age for females was 54.3 ± 18.8 years, and for males 55.4 ± 18.3 years. The dataset covers a wide range of adult lifespan and equitable gender representation. The Cam-CAN dataset included data acquired with consistent scanning protocols and quality control standards, making it a reliable source for training models and exploring brain aging. Detailed MRI data acquisition protocol can be found in the Supplementary Material S1 Appendix and Cam-CAN’s website (https://camcan-archive.mrc-cbu.cam.ac.uk).

The preprocessing of fMRI data was conducted in SPM12, automated and parallelised by Automatic Analysis (aa) [74] pipeline, which included the following major steps: (1) slice timing correction, (2) head motion and distortion correction, (3) spatial normalization and smoothing, (4) interference signal removal, (5) band-pass filtering, and region of interest (ROI) time series extraction with the standard Automated Anatomical Labeling (AAL) [75] atlas (Listed in Supplementary Material S1. Table). Pearson correlations between all pairs of regional blood oxygen level-dependent signals were calculated for each participant, and the whole-brain functional network (90 × 90) was constructed. For the sMRI data, preprocessing was performed using DARTEL [76]. Then, each volumetric image was spatially resampled to a fixed resolution of 128×128×128 voxels. Following resampling, intensity normalization was applied by scaling voxel intensities to the range [0, 1], enhancing the numerical stability of the training process and facilitating convergence.

Additionally, this work further evaluated the proposed method using neuroimaging data from the Open Access Series of Imaging Studies, Phase 3 (OASIS-3) [77]. Specifically, we evaluated the developed method with T1 structural and resting-state functional images from 533 cognitively normal participants (236 males, 297 females) with a mean age of 68.9 ± 8.7 years and an age range of 42.0–95.0 years. The mean age of females was 67.9 ± 9.1 years, while males had a mean age of 70.2 ± 8.1 years. Detailed information about the dataset and imaging protocols is available on the OASIS website (https://sites.wustl.edu/oasisbrains/home/oasis-3/). The data protocol of OASIS-3 and its preprocessing can be seen in Supplementary Material S2 Appendix.

### fMRI branch

As shown in Figure 1, the fMRI branch includes two modules: (1) Whole brain Graph Construction: For each participant, the preprocessed ROI-wise time series are used to compute a whole-brain FC matrix based on pair-wise Pearson cross-correlations. This matrix is then transformed into a graph representation, where each node represents a brain ROI and each edge weight represents the FC between regions. (2) Feature Representation by Graph Attention Network (GAT): A multi-layer GAT is employed to learn informative representations from the constructed graphs. The attention mechanism enables the model to extract node-level features that capture region-specific activation patterns and assign adaptive weights to edges, thereby identifying important functional connectivities for prediction.

**Fig 1.**
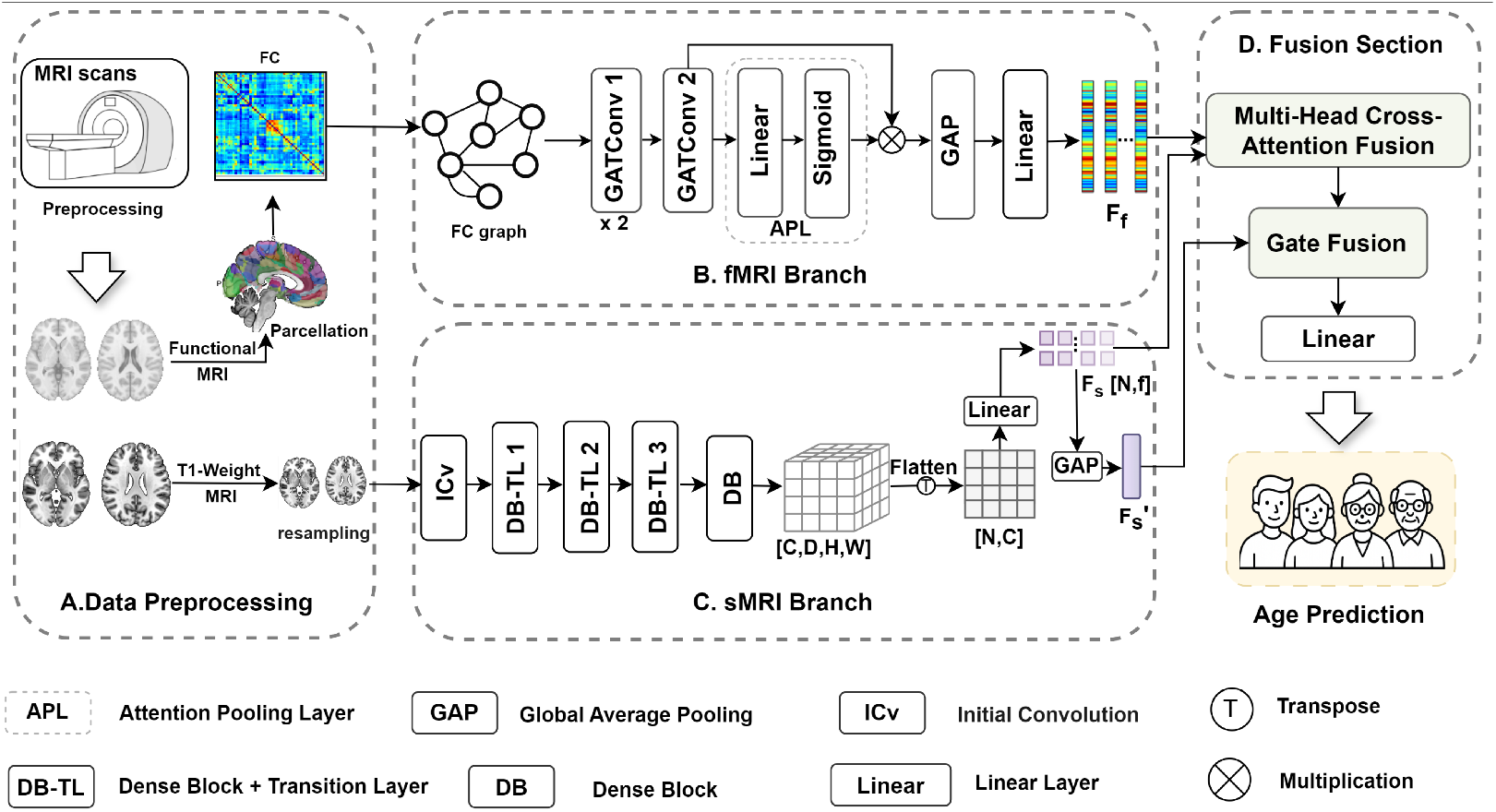
Overview of the Attention-guided Multimodal brain Age prediction Net (AMAge-Net) framework. **A**. Preprocessing steps of sMRI and rs-fMRI data. The original FC matrices are obtained from the fMRI time series data with the AAL atlas. **B**. The fMRI branch.Construction of individual brain graphs based on FC matrices and representation learning by a multi-layer Graph Attention Network (GAT). **C**. The sMRI branch. A 3D DenseNet121-based transformation is implemented on the preprocessed T1w images to capture voxel-level hierarchical and multi-scale structural features through dense connections, resulting in structural feature representations. **D**. Fusion section. A Multi-Head Cross-Attention mechanism followed by a Gated Fusion module is adopted for cross-modal integration, and a fully connected layer is used to make the final prediction.

#### Whole brain Graph Construction

We construct a node feature graph *G* = (*V, E*) based on the FC matrix to transform FC data into a graph-structured representation. Brain ROIs are represented as graph nodes, defined as *V* = {*v*_1_, …, *v*_*n*_*}*, and the set of edges is denoted as *E*. Edges are the strength of functional connectivities, and node features are derived from each region’s connectivity profile in the graph. The construction process is detailed as follows:

1. Nodes: Each brain ROI corresponds to a node in the graph, resulting in a total of *n* nodes (*n*=90).
2. Node feature: The feature vector of each node is defined as its connectivity strength with all other nodes based on the FC matrix. All node feature vectors form the node feature set *X* = {*X*_1_, *X*_2_, …*Xn*}, *X*_*i*_ ∈ *R*^*d*^, where *X*_*i*_ is the feature vector associated with node *v*_*i*_, *d* denotes the initial feature dimension.
3. Edge: An edge between two nodes is defined as the absolute value of their functional connectivity, |*FC*_*ij*_|, where *FC*_*ij*_ denotes the PCC between node *v*_*i*_ and node *v*_*j*_; specifically, an edge is present if |*FC*_*ij*_| ≠ 0, in which case |*FC*_*ij*_| is assigned as the edge weight.

The additional details on the construction of the whole-brain functional graph can be seen in S1 Appendix.

#### Feature Representation by Graph Attention Network

The feature extraction from fMRI begins with the input graph *G* and node features *X*. The first graph attention layer applies Multi-Head Attention(MHA) with H heads [53], projecting each node’s feature vector *X*_*i*_ into a hidden space *d*^′^. For each attention head h, attention coefficients between connected nodes *v*_*i*_ and *v*_*j*_ are computed as:

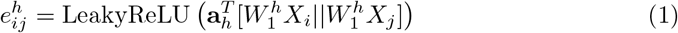

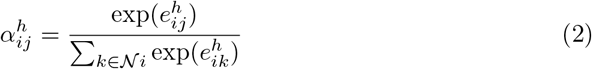

where *W*_1_ is a learnable weight matrix and *a*_*h*_ is the attention vector for head h. *X*_*i*_ represents the feature vector for the *i*-th node in the graph. *X*_*j*_ represents the feature vector of a neighboring node *v*_*j*_ that is connected to node *v*_*i*_ (*v*_*j*_ ∈ 𝒩_*i*_, 𝒩_*i*_ denoted as the set of neighboring nodes of node *v*_*i*_. The output node representation of the layer is computed as:

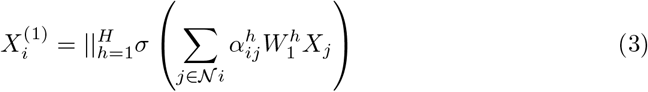

The second graph attention layer takes *X*^(1)^ as input and repeats the same MHA process, producing a refined representation:

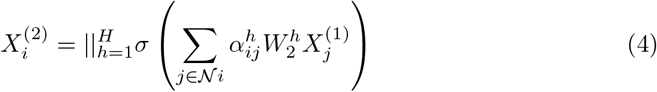

A third graph attention layer is applied with a single attention head and without concatenation, reducing the feature dimension back to *d*^′^ (the hidden feature space):

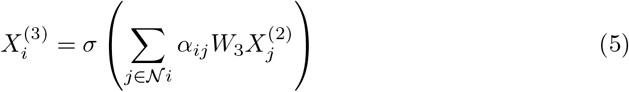

Then, an attention pooling mechanism is applied to obtain the graph-level feature representation:

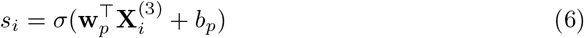

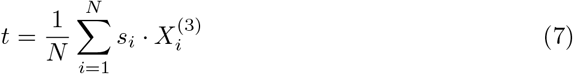

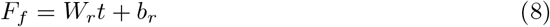

Where *s*_*i*_ represents the importance score of each node, *σ* is the sigmoid activation function and *w*_*p*_, *b*_*p*_ and *b*_*r*_ are learnable parameters. *W*_*r*_ is the learnable weight matrix associated with the h-th attention head, where *W*_*r*_ ∈ ℝ^*k*×*d*^*′* projects the graph representation *F*_*f*_ to the output dimension *k*.

### sMRI branch

In the sMRI branch, as shown in Figure 1 and Figure 2, the T1w image is resampled and fed into a convolutional neural network based on 3D DenseNet121 to extract deep spatial features. The network begins with an initial convolutional module consisting of a 3D convolutional layer, batch normalization, a LeakyReLU activation function, and 3D max pooling, which is used to extract the initial feature maps. Subsequently, the features are passed through four Dense Blocks, interleaved with three Transition Layers. The four Dense Blocks contain 6, 12, 24, and 16 Bottleneck units, respectively. Each Bottleneck unit consists of batch normalization, a LeakyReLU activation function, and convolutional layers. Each Transition Layer includes batch normalization, a LeakyReLU activation function, a convolutional layer, and max pooling.

**Fig 2.**
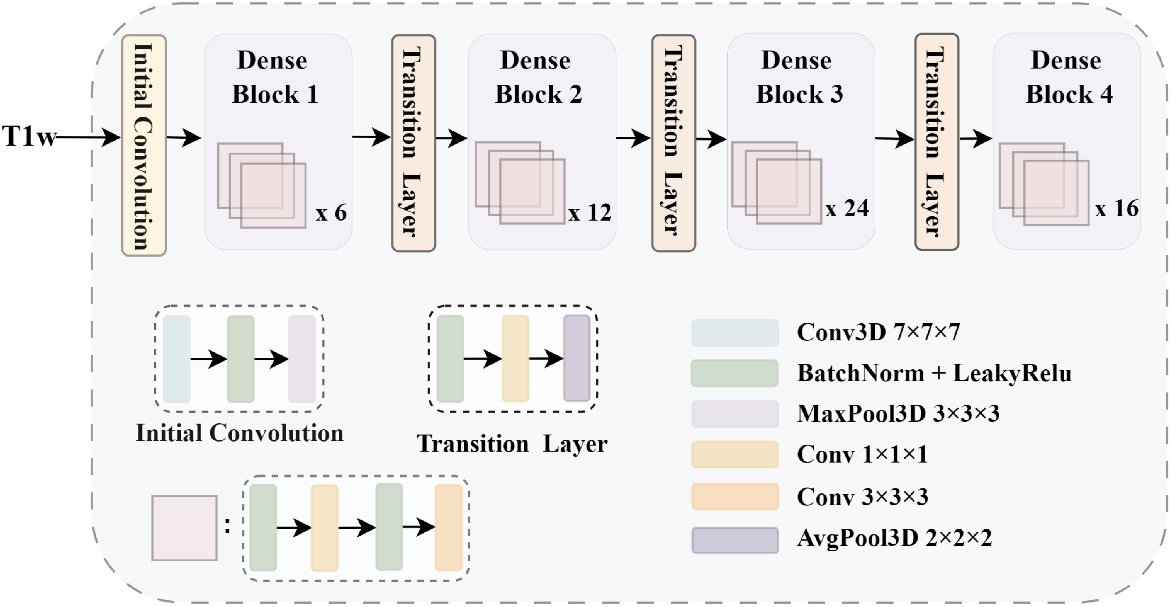
Feature extraction from sMRI using 3D-DenseNet121.

The structural feature map output from the fourth Dense Block is flattened and projected to a lower-dimensional space to obtain the structural feature representation 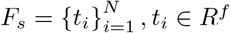,where *N* is the number of spatial locations and *f* is the feature dimension. This representation is subsequently used for feature fusion. In addition, a global average pooling operation is applied to *F*_*s*_ to generate a global structural representation vector 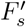, which captures the overall anatomical characteristics of each participant.

### Multimodal Data Fusion and Age Prediction

To effectively integrate complementary information from functional and structural modalities, we design a two-stage fusion strategy, as shown in Figure 3. First, we employ a Multi-Head Cross-Attention module to align and fuse the structural feature representation with the fMRI-derived feature representation, enabling fine-grained interaction between anatomical features and functional connectivity patterns. Then, to further enhance the integration, we introduce a Gated Fusion module that adaptively combines the global structural representation with the feature representation from the first stage. This hierarchical fusion strategy allows the model to incorporate both local anatomical details and global structural context into the final representation, thereby improving the model’s ability to capture multi-level brain characteristics for brain age prediction.

**Fig 3.**
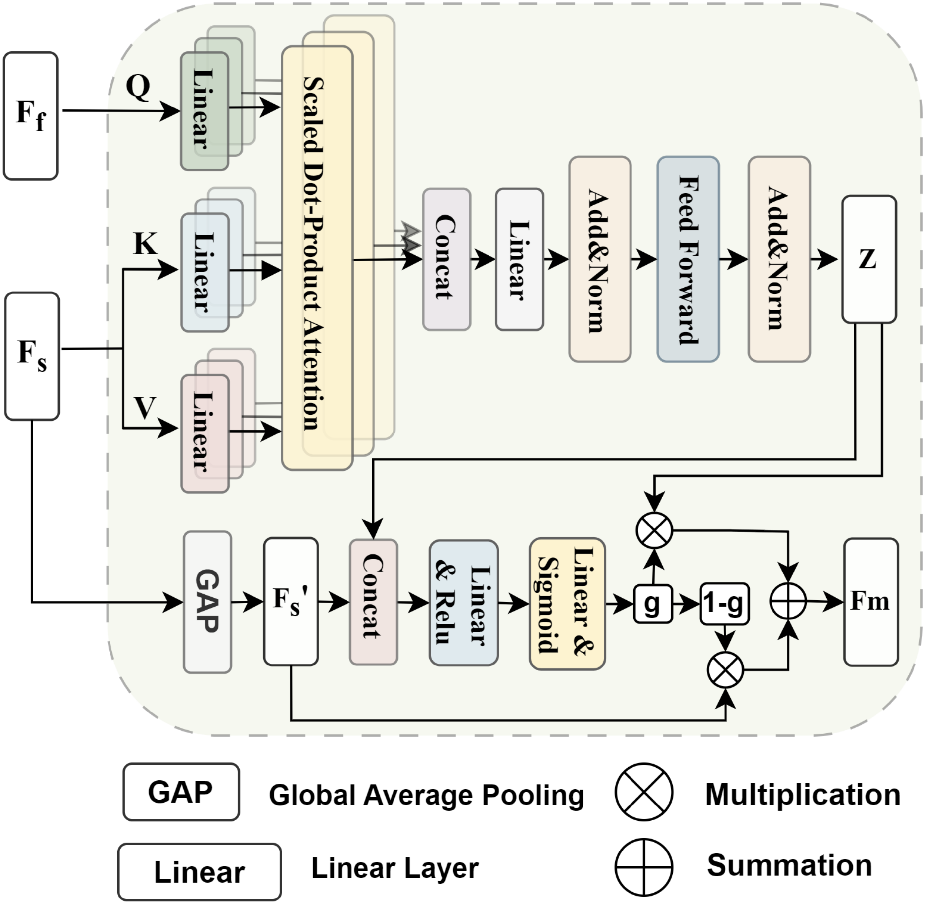
Fusion Section. The inputs are functional features *F*_*f*_ and structural features *F*_*s*_. A Multi-Head Cross-Attention mechanism produces the fused vector *Z*. Then, the final fused feature *F*_*m*_ is obtained through Gated Fusion, where *g* is the gate weight.

#### Multi-Head Cross-Attention Fusion

A Multi-Head Cross-Attention Fusion mechanism is adopted to integrate structural and functional features with two inputs: a functional feature vector *F*_*f*_ and a structural feature representation *F*_*s*_. In this mechanism, *F*_*f*_ serves as the guidance for selectively aggregating information from *F*_*s*_. Then, linear layers are used to generate the query, key, and value representations with these inputs separately. Specifically, *F*_*f*_ is transformed into the query, while *F*_*s*_ is transformed into the key and value vectors. These transformed representations are then fed into multiple scaled dot-product attention heads. Each attention head calculates attention scores by comparing the query with the key, scales the scores, applies a softmax function, and then uses these weights to aggregate the value vectors. The outputs from all attention heads are then concatenated and passed through a linear layer. The result is subsequently processed by a residual feed-forward module with layer normalization, yielding the final fused output, denoted as *Z*. This multi-head structure enables the model to capture diverse contextual relationships within the input data in parallel, improving its capacity to focus on different parts of the input simultaneously and effectively learn complex dependencies.

Given the structural feature representation *F*_*s*_ ∈ *R*^*N*×*f*^ and the functional feature vector *F*_*f*_ ∈ *R*^*k*^, where *f* = *k*, the query, key, and value matrices are obtained through linear projections:

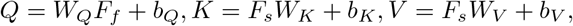

where *W*_*Q*_, *W*_*K*_, *W*_*V*_ and *b*_*Q*_, *b*_*K*_, *b*_*V*_ are learnable parameters. Then we compute scaled dot-product attention for each head [53]:

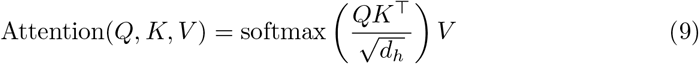

where *d*_*h*_ = *f/m* is the dimensionality of each attention head and *m* is the number of heads. The outputs from all heads are concatenated and linearly projected to form the fused representation:

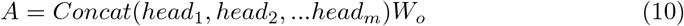

Finally, the fused feature *A* is passed through a feed-forward network followed by layer normalization to obtain the final multimodal representation *Z*. By allowing information exchange across modalities through multiple attention heads, the Multi-Head Cross-Fusion module explicitly models fine-grained interactions between structural and functional features at the regional level and enables the model to capture complementary and localized cross-modal relationships that are important for age prediction.

#### Gated Fusion

To further integrate hierarchical multimodal information and enhance predictive performance, we introduce a Gated Fusion mechanism that adaptively combines the Multi-Head Cross-Attention derived features *Z* with the global structural representation vector 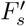. We concatenate the feature *Z* with 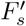 to form the fused input feature 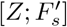. A linear transformation followed by a sigmoid activation function is applied to the concatenated vector to compute the fusion weight *g*:

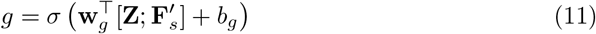

where *w*_*g*_ and *b*_*g*_ are learnable parameters.The final fused feature is

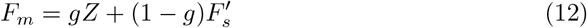

The Gated Fusion module operates at a higher level to adaptively integrate the cross-fused features with global structural representations. By learning modality-dependent weights, it allows the model to dynamically balance local cross-modal information and holistic anatomical context across subjects, improving robustness and preventing over-reliance on a single modality.

#### Regression Prediction

In this study, the regression task aims to predict the brain age of each participant based on multimodal neuroimaging features. The final feature representations are obtained through the fusion module that integrates sMRI and fMRI information. The fused feature *F*_*m*_ is fed into a fully connected regression layer to produce a scalar output representing the predicted brain age. The model is optimized using the Mean Squared Error (MSE) loss function, which penalizes large deviations between predicted brain age and the chronological age (ground truth). This setup enables the model to learn a continuous mapping from integrated brain features to the corresponding age value.

### Performance Evaluation

#### Model Evaluation Metrics

In this study, a five-fold cross-validation strategy was used to evaluate the performance of different methods. Specifically, the participants were divided into five approximately equal-sized subsets, with one subset as the validation set and the remaining four as the training sets in each iteration. The result presented in this paper is the average over five subsets. Model performance was comprehensively evaluated using four metrics: Mean Absolute Error (MAE), Root Mean Square Error (RMSE), R-squared (*R*^2^), and Pearson cross-correlations (PCC). Specifically, MAE measured the average absolute difference between predicted brain age and actual chronological age, reflecting prediction accuracy in years; RMSE computed the square root of the prediction errors, penalizing larger errors more heavily; *R*^2^ indicated the proportion of variance in actual age explained by the model; and PCC quantified the linear correlation between predicted and actual values. The details of the Evaluation Metrics can be seen in the Supplementary Material S1 Appendix. Additionally, the contribution of individual brain regions to the prediction was analyzed by modality-specific saliency measures. To be specific, region-wise importance was derived from the learned attention weights within the hierarchical graph attention network for the fMRI branch, and gradient-based saliency maps were utilized to calculate importance scores of brain regions for the sMRI branch [11]. These help to reveal neuroanatomical features associated with aging and enhance the interpretability of the model.

#### Comparison with SOTA

We compared our proposed method with other methods recently published in the literature [30, 39–41, 50, 52, 58]. The compared methods are based on deep neural networks, including the GNN [30, 50, 52], CNN [39–41], and transformer [58]. For a fair comparison, these competing methods were trained using learning schemes (see Supplementary Material S1 Appendix for more details about the implementation setup as well as computational cost) that are highly consistent with those used in our method, as detailed below.

- **BrainGNN**: Kumar et al. [30] presents a GNN-based framework for brain age prediction at the ROI level. We modeled each participant’s brain as a graph, where each ROI is represented as a node to learn ROI-based graph embeddings. Node features are first embedded into a low-dimensional latent space. Subsequently, graph pooling or coarsening operations are applied to reduce and aggregate the node representations. The resulting summary vector is then fed into a Multi-Layer Perceptron (MLP). The entire framework—including the graph convolution, pooling layers, and the MLP—is trained in an end-to-end manner to automatically learn discriminative features from brain graphs.
- **GCN**: We use graphs constructed from FC matrices as inputs to the GCN [50]. The GCN consists of two graph convolution layers, a global pooling layer, and two fully connected layers, with the final layer outputting the predicted age.
- **BC GCN**: This method is proposed by Li et al. [52]. It introduces an edge-based Graph Path Convolution (GPC) framework that aggregates information from multiple graph paths to capture high-order structural dependencies. To reduce the graph size while preserving key topological features, it employs both Edge Pooling (E-P) and Node Pooling (N-P) strategies for hierarchical downsampling. Furthermore, Residual blocks [78] and Squeeze-and-Excitation (SE) blocks [79] are incorporated alongside the GPC layers to enhance feature representation and facilitate brain age prediction.
- **ResNet**: Ning et al. [39] used the ResNet model provided by NiftyNet [78] for brain age prediction. The model consists of one convolutional layer, six bottleneck layers, one fully connected layer, and an output layer. Multiple shortcut connections exist between the bottleneck layers. The initial convolutional layer contains 64 filters. There are six bottleneck layers in total: bottleneck layers 1 and 2 have 128 filters; bottleneck layers 3 and 4 have 256 filters; and bottleneck layers 5 and 6 have 512 filters.
- **EfficientNet**: Poloni et al. [40] adopted a 3D-adapted EfficientNet architecture for brain age prediction. The network consists of 16 mobile inverted bottleneck convolution (MBConv) blocks, with all original 2D operations replaced by 3D operations. After processing the input image, the final output layer uses a linear activation function for age regression.
- **3D-CNN**: Hong et al. [41] proposed a 3D CNN architecture to predict brain age from conventional brain magnetic resonance images in children. The model includes seven 3D convolutional layers and three fully connected layers. We feed the resampled MR images into this model to perform brain age prediction.
- **MFFormer**: Wang et al. [58] proposed method adopts a dual-branch architecture: one branch processes 2D rs-fMRI time series, while the other processes 3D T1w MRI. A Transformer fusion module is then used to integrate the features. After aligning the multi-dimensional features using a dimension-up/dimension-down strategy, self-attention is applied to capture cross-modal nonlinear dependencies. Finally, a lightweight classification head (consisting of 2 convolutional layers and 3 MLP layers) outputs the diagnostic result. In our work, we modified the final classification head into a regression layer for brain age prediction.
- **CTransformer**: Zhang et al. [59] proposed a Transformer-based multimodal framework that integrates structural and functional MRI using a dual-branch architecture. The structural branch employs a 3D CNN with Transformer Attention to capture global anatomical patterns, while the functional branch uses a cascaded Transformer to encode functional connectivity relationships. Cross-attention–based fusion is applied to integrate functional information into hierarchical structural representations. In our work, we replace the original classification head with a regression layer to enable brain age prediction.

#### Association Analysis and gender differences

To further validate the relationship between brain FC and chronological age, we performed a whole-brain correlation analysis between FC and age. To control for false positives due to multiple comparisons, false discovery rate (FDR) correction was applied to the correlation results to ensure statistical reliability. Following this, the brain regions identified from the correlation analysis were compared with the important brain regions derived from the brain age prediction model, providing a complementary evaluation of model performance and its neurobiological interpretability.

To investigate sex-related differences in brain age prediction, we trained and evaluated the same model architecture separately for male (n = 322) and female (n = 330) groups using the same hyper-parameters and training strategy. This stratified approach allowed us to assess whether the brain regions contributing to age prediction differ between sexes. After training, the most influential brain regions contributing to prediction were identified separately for the male and female groups, and a ranked analysis of their importance was conducted.

## Results

### Brain Age Prediction Performance

We report the experimental results of our proposed AMAge-Net and nine competing methods in brain age prediction in Table 1. In Figure 4, we present a comparative analysis based on scatter plots illustrating the relationship between predicted and true age across eight different methods. Two key evaluation metrics—MAE and *r* —are used to assess the accuracy and consistency of predictions. From Table 1 and Figure 4, we have the following findings.

**Table 1.** Result comparison of our proposed method with other methods in brain age estimation.

| Model | Data Modality | MAE | RMSE | R<sup>2</sup> | PCC (r) |
| --- | --- | --- | --- | --- | --- |
| BrainGNN [30] | fMRI | 11.46 | 14.55 | 0.37 | 0.62 |
| GCN [50] | fMRI | 12.99 | 15.97 | 0.25 | 0.51 |
| BC_GCN [52] | fMRI | 8.39 | 10.52 | 0.67 | 0.82 |
| ResNet [39] | sMRI | 7.34 | 9.29 | 0.74 | 0.86 |
| EfficientNet [40] | sMRI | 12.80 | 15.18 | 0.32 | 0.57 |
| 3D-CNN [41] | sMRI | 5.76 | 7.39 | 0.84 | 0.92 |
| MFFormer [58] | fMRI+sMRI | 8.13 | 10.22 | 0.69 | 0.84 |
| CTransformer [59] | fMRI+sMRI | 8.40 | 10.93 | 0.65 | 0.81 |
| AMAge-Net-Concat | fMRI+sMRI | 5.29 | 6.77 | 0.86 | 0.93 |
| **AMAge-Net (Ours)** | fMRI+sMRI | **5.09** | **6.52** | **0.87** | **0.94** |

**Fig 4.**
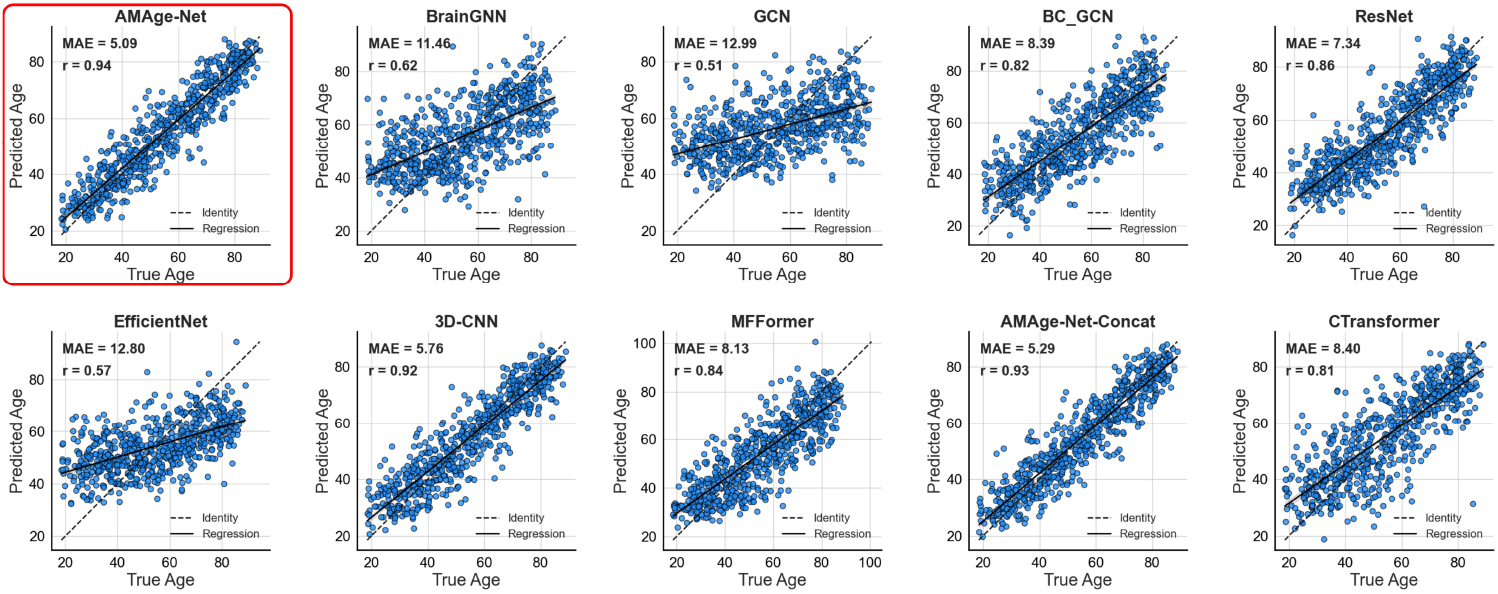
The scatter plots of predicted versus chronological age for AMAge-Net (see red box), three multimodal models, and six unimodal models, demonstrating the accuracy of brain age estimation.

First, our model significantly outperforms all baseline methods across all evaluation metrics on the main dataset. In particular, it achieves the lowest MAE = 5.09, lowest RMSE = 6.52, highest *R*^2^ = 0.87, and the highest r = 0.94. Compared to the best-performing single-modality baseline, our model improves MAE by 11.6% and PCC by 2%. This demonstrates that the proposed model can better capture age-related brain features by leveraging both structural and functional information through attention-based fusion. Second, models using only functional features generally perform worse than structural features or multimodal approaches. For instance, BrainGNN yields an MAE of 11.46 and an *R*^2^ of only 0.62, which highlights the limited predictive power of functional features alone for brain age estimation. In contrast, methods that utilize structural features (such as 3D-CNN and ResNet) show notably better results, suggesting that sMRI provides stronger age-related signals. Third, although multimodal fusion is generally expected to enhance performance by integrating complementary information from structural and functional modalities, not all fusion strategies effectively leverage this potential.

Among the multimodal methods considered, we implemented a direct feature concatenation–based fusion strategy on top of the proposed dual-branch architecture as a strong baseline (denoted as AMAge-Net-Concat), whereas MFFormer and CTransformer follow the fusion designs reported in their original studies [58, 59]. Although all methods integrate structural and functional information, MFFormer (MAE = 8.13, PCC = 0.84) and CTransformer (MAE = 8.40, PCC = 0.81) underperform compared with the direct concatenation baseline AMAge-Net-Concat, in our experiments (MAE = 5.29, PCC = 0.93). In contrast, our proposed model incorporates a Multi-Head Cross-Attention mechanism and a Gated Fusion strategy, enabling a more dynamic and adaptive integration of structural and functional features. As a result, the proposed AMAge-Net further improves upon direct concatenation, achieving gains of 3.8% in MAE and 1.1% in PCC. These results indicate that effective multimodal learning depends not only on the inclusion of multiple modalities, but also critically on the fusion mechanism’s ability to capture complementary cross-modal information.

We further validated the model on an independent external OASIS-3 dataset with 533 subjects (see Methods for details). The model achieves an MAE of 4.29, RMSE of 5.59, *R*^2^ of 0.58, and PCC of 0.77 (as shown in Supplementary Material S1. Figure), demonstrating that it maintains strong predictive performance across different cohorts and consistently outperforms both single-modality and existing multimodal baselines (Supplementary Material S2. Table). This highlights the robustness and generalizability of our approach.

### Identifying Multimodal Brain Patterns of Aging

To further interpret the proposed model and identify neuroanatomical correlates of brain aging, we analyzed the contribution of individual brain regions to the prediction using modality-specific saliency measures (denoted as *S*_*fMRI*_ and *S*_*sMRI*_, respectively) S3. Table and S4. Table list the specific contribution for each region from fMRI and sMRI.

For the fMRI branch, region-wise importance was derived from the learned attention weights within the hierarchical GAT, highlighting functional nodes most influential for age estimation. We retained the top 24 regions that contribute most to age prediction performance, as indicated by the highest saliency scores for each subject (score *>* 0.7). This threshold provides a compact yet informative functional representation while maintaining a fixed feature dimensionality across individuals. It balances information preservation and noise reduction, and the identified regions consistently span multiple large-scale functional networks associated with aging-related functional changes. The top contributing brain regions are shown in Figure 5, and also listed in Supplementary Material S5. Table. These top-ranked regions included the right thalamus (*S*_*fMRI*_ = 1), left superior medial frontal gyrus (*S*_*fMRI*_ = 0.99), left insula(*S*_*fMRI*_ = 0.94), right middle temporal pole (*S*_*fMRI*_ = 0.92), and left caudate (*S*_*fMRI*_ = 0.88), along with several prefrontal and cingulate regions. Besides, the functional connectivity links involving these saliency brain regions, including the inferior temporal gyrus, the superior temporal pole, and the postcentral gyrus, were found to be significantly correlated with age in the association analysis. A list of the top 200 significantly related functional connectivities can be found in S5. Table.

**Fig 5.**
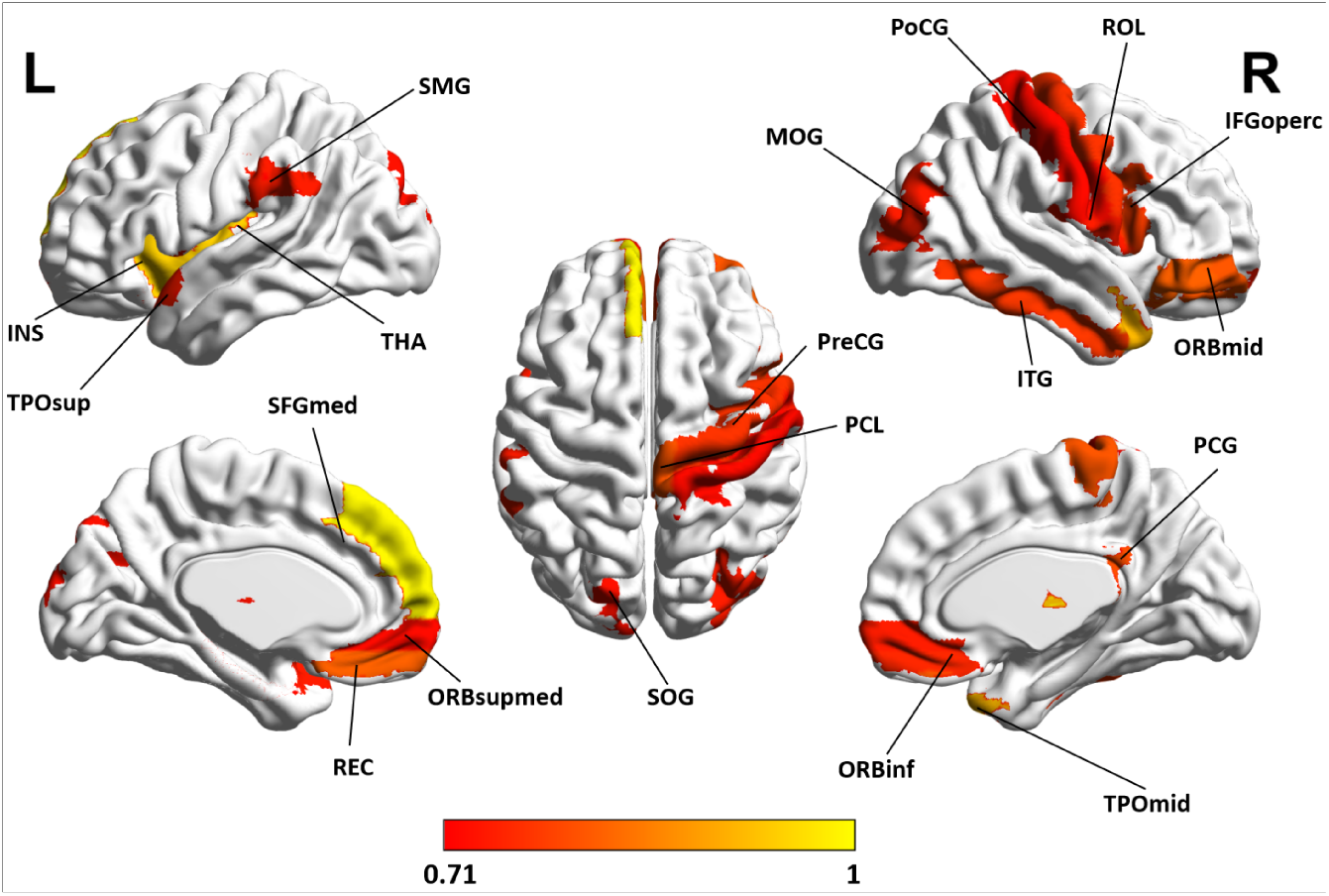
Distribution of brain region importance derived from fMRI data. Region-level importance scores were computed from the node-wise attention weights learned by the graph attention network and subsequently normalized to the range [0,1]. To emphasize the most influential functional regions, only brain regions with normalized importance scores greater than 0.7 are displayed.

For the sMRI branch, importance scores were computed from gradient-based saliency maps on the 3D DenseNet121 features. In order to maintain the same number of selected regions as in the fMRI branch, the top contributing brain regions with an importance score over 0.5 (23 regions) are shown in Figure 6 and listed in S6. Table, revealing a predominance of posterior and subcortical structures such as the right posterior cingulate (*S*_*sMRI*_ = 1.0), right precuneus (*S*_*sMRI*_ = 0.83), right calcarine (*S*_*sMRI*_ = 0.728), left posterior cingulate (*S*_*sMRI*_ = 0.725), and right cuneus (*S*_*sMRI*_ = 0.716). Notably, several regions (e.g., right thalamus, right paracentral lobule, right posterior cingulate) were salient in both modalities, suggesting convergent functional–structural markers of brain aging.

**Fig 6.**
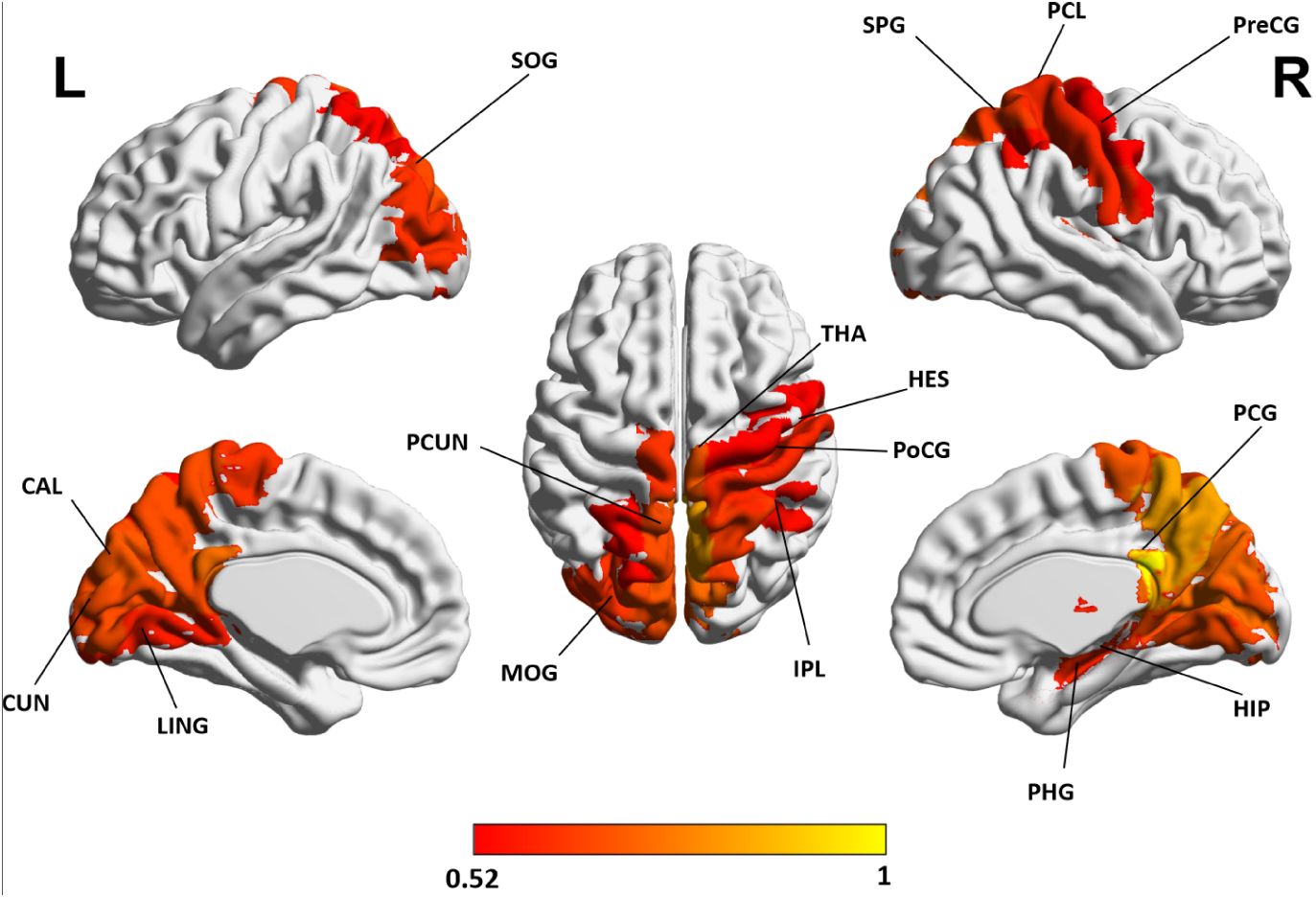
Distribution of brain region importance derived from sMRI data. Region-level importance scores were computed using gradient-based saliency analysis, in which voxel-wise saliency values were aggregated within each brain region and subsequently normalized to the range [0,1]. To emphasize the most influential regions, only brain regions with normalized importance scores greater than 0.5 are displayed.

In addition, we applied the same prediction model separately to male and female groups to examine sex-specific brain-aging patterns. To ensure comparability with the overall model without gender stratification, we selected the top 23 contributing regions for the sMRI branch and the top 24 contributing regions for the fMRI branch, matching the respective numbers used in the overall analysis. Similarity between male and female saliency maps was quantified as the percentage of overlapping brain regions between the two sets of top-ranked regions. For the sMRI branch, the top 23 contributing regions showed strong concordance between sexes (similarity *>* 70%), notably the posterior cingulate cortex, precuneus, hippocampus, and thalamus (Figure 8, S6. Table). By contrast, the fMRI branch exhibited greater sex-dependent divergence with similarity around 20% of the top 24 contributing regions (Figure 7, Table S7. Table); only a handful of regions were shared, mainly within the frontal lobe (inferior orbitofrontal gyrus, inferior opercular frontal gyrus, and olfactory cortex). Notably, the motor cortex emerged as an important contributor only in males, whereas fusiform and occipital regions were prominent only in females.

**Fig 7.**
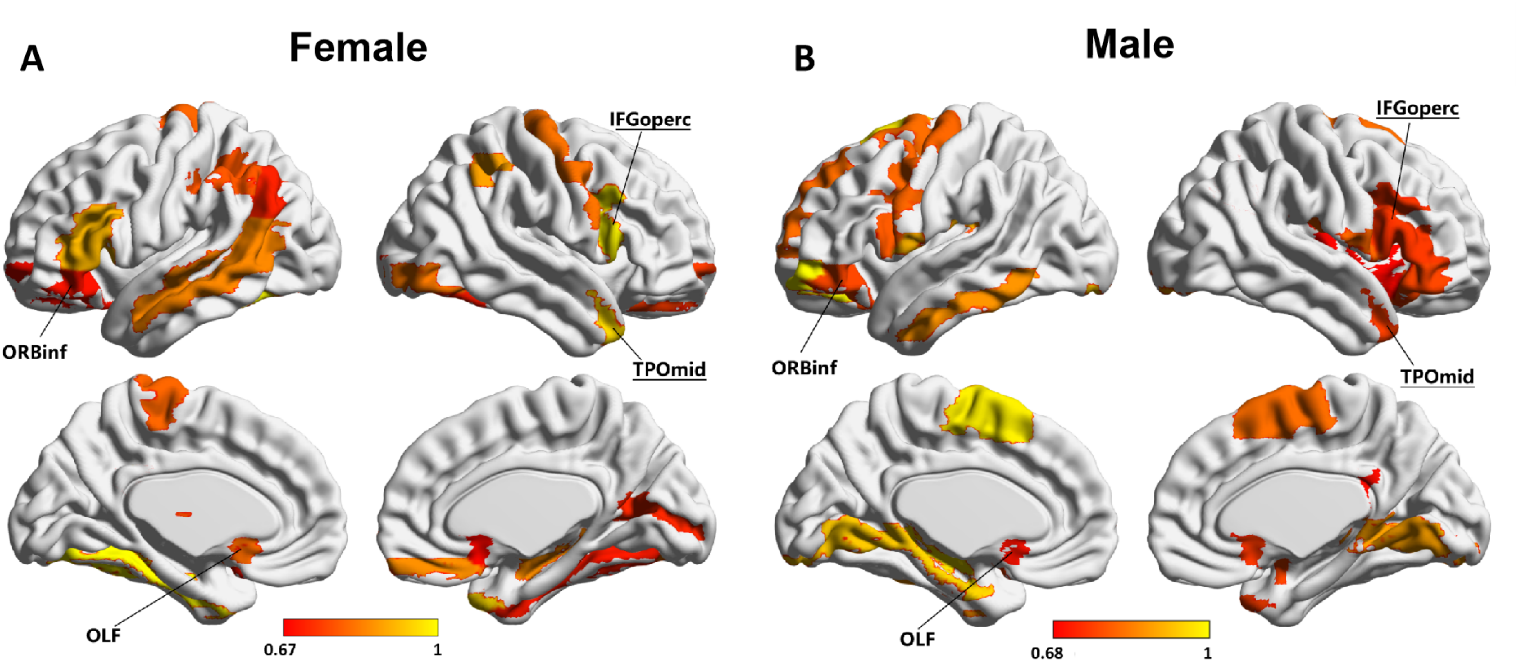
Distribution of brain region importance derived from attention weights computed on fMRI data in the sex-specific brain age prediction analysis. The top 24 contributing regions are shown to match the respective numbers used in the overall analysis. (A) Results obtained using only female participants. (B) Results obtained using only male participants.

**Fig 8.**
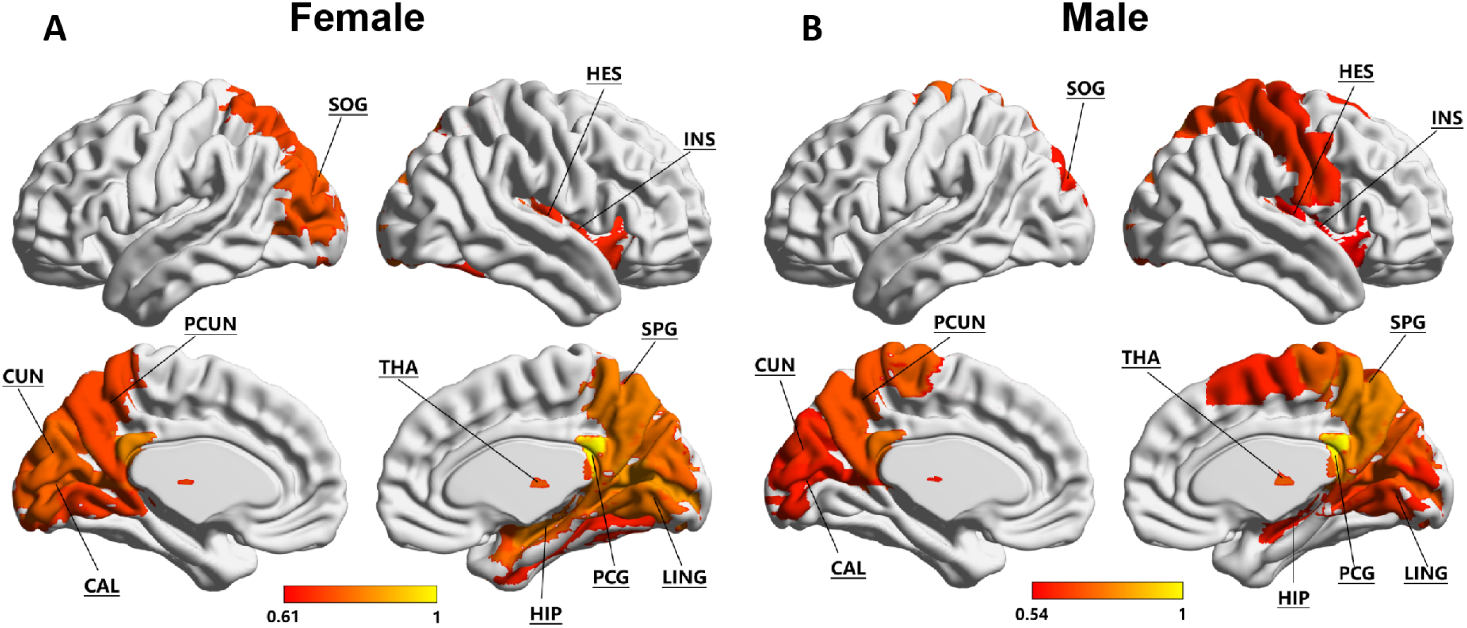
Distribution of brain region importance derived by saliency maps computed from sMRI data in the sex-specific brain age prediction analysis. The top 23 contributing regions are shown to match the respective numbers used in the overall analysis. (A) Results obtained using only female participants. (B) Results obtained using only male participants.

### Ablation Study

To evaluate the individual contributions of each component within our proposed framework, in the Table 2 and Figure 9, we conducted a series of ablation experiments using the same dataset. Specifically, we compared the full AMAge-Net model against four variants:

**Table 2.**
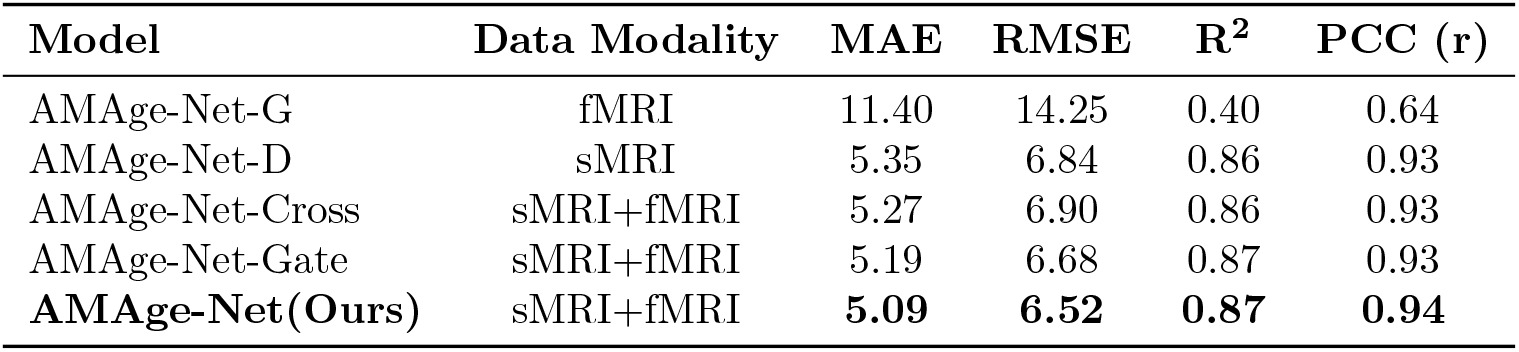
Ablation studies compare different models and feature combinations. Evaluations include baseline models using only structural or functional features, models using only Cross-fusion or only Gated fusion, as well as our proposed method.

| Model | Data Modality | MAE | RMSE | $R^2$ | PCC (r) |
| --- | --- | --- | --- | --- | --- |
| AMAge-Net-G | fMRI | 11.40 | 14.25 | 0.40 | 0.64 |
| AMAge-Net-D | sMRI | 5.35 | 6.84 | 0.86 | 0.93 |
| AMAge-Net-Cross | sMRI+fMRI | 5.27 | 6.90 | 0.86 | 0.93 |
| AMAge-Net-Gate | sMRI+fMRI | 5.19 | 6.68 | 0.87 | 0.93 |
| **AMAge-Net(Ours)** | sMRI+fMRI | **5.09** | **6.52** | **0.87** | **0.94** |

**Fig 9.**
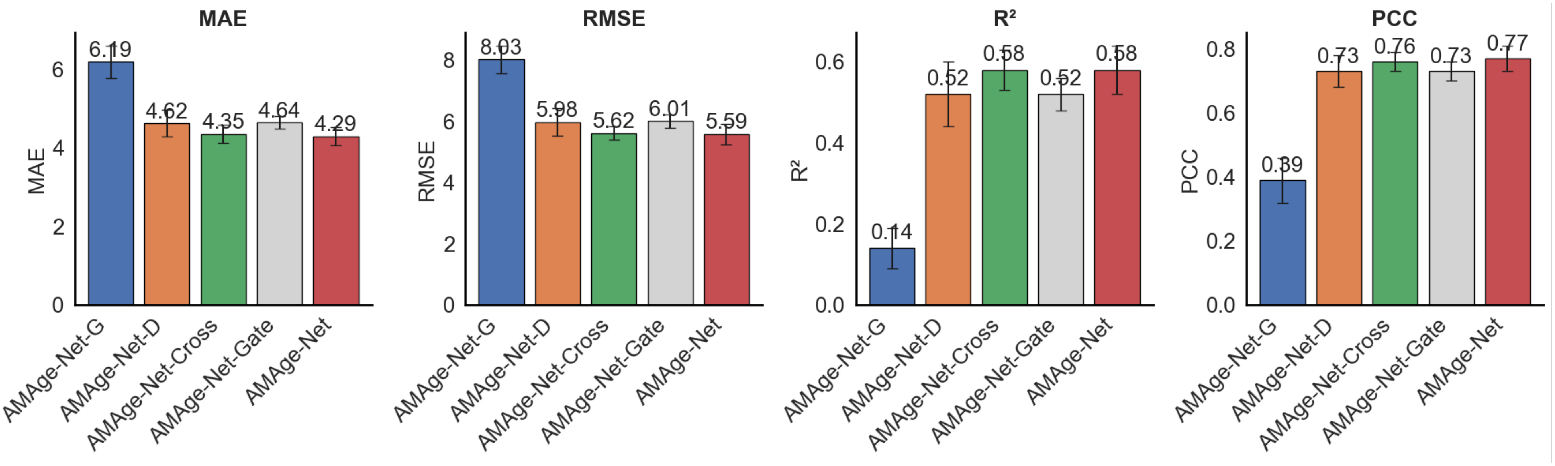
Ablation study across different models. The performance metrics include MAE, RMSE, *R*^2^, and PCC.

- **AMAge-Net-G**, which uses only **fMRI** features processed by GAT in the isolated functional branch (see Figure 1B for the sMRI branch), without structural input.
- **AMAge-Net-D**, which uses only **sMRI**features extracted by the DenseNet121 in the isolated structural branch (see Figure 1C for the sMRI branch), without incorporating functional information.
- **AMAge-Net-Cross**, which performs multimodal fusion using only the **Cross-Fusion** module, enabling cross-modal interactions while excluding gated feature selection.
- **AMAge-Net-Gate**, which performs feature fusion using only the **Gated Fusion** module, focusing on adaptive feature weighting without explicit cross-modal attention.

The ablation study was designed to disentangle the individual contributions of the sMRI and fMRI branches and the two complementary fusion mechanisms, where single-modality variants assess structural and functional information independently, and the Cross-Fusion and Gated Fusion variants isolate explicit cross-modal interaction and adaptive feature weighting, respectively, thereby justifying the architectural design of AMAge-Net.

We first evaluated the baseline performance of using only the functional features processed by GAT in the functional branch, AMAge-Net-G. This model achieved a MAE of 11.40 and a RMSE of 14.25, with an *R*^2^ of 0.40 and a PCC of 0.93. In contrast, when using only structural features, DenseNet121 in the isolated structural branch, performance increases significantly (MAE = 5.35, RMSE = 6.84, *R*^2^ = 0.86, PCC = 0.93), highlighting the relatively limited predictive power of isolated structural information.

Next, we examined the contributions of different fusion strategies within the multimodal framework. AMAge-Net-Cross, which uses only the Multi-Head Cross-Attention module, explicitly models interactions between structural and functional features and outperforms unimodal models, demonstrating the benefit of capturing fine-grained cross-modal relationships for brain age estimation (MAE = 5.27, RMSE = 6.90, *R*^2^ = 0.86, PCC = 0.93). In contrast, AMAge-Net-Gate relies solely on the Gated Fusion mechanism to adaptively weight multimodal features without explicit cross-modal attention, resulting in comparatively limited performance (MAE = 5.19, RMSE = 6.68, *R*^2^ = 0.87, PCC = 0.93).

The full AMAge-Net, which combines Multi-Head Cross-Attention with Gated Fusion, achieves the best overall performance (MAE = 5.09, RMSE = 6.52, *R*^2^ = 0.87, PCC = 0.94). The cross-attention module captures fine-grained ROI-level interactions between modalities, while the gated fusion module adaptively integrates these cross-fused features with global structural representations. Compared with single-modality models, the full model reduces MAE by 1.33 and 0.60, and RMSE by 1.43 and 0.63, for fMRI-only and sMRI-only inputs, respectively. These results highlight the effectiveness of jointly modeling explicit cross-modal interactions and adaptive feature weighting. Importantly, we conducted the same ablation study on the OASIS-3 dataset and observed consistent performance trends across all variants (see Supplementary Material S2. Figure and S8. Table), further supporting the robustness of the proposed fusion design.

## Discussion

Accurate estimation of brain age from neuroimaging data has emerged as a promising biomarker for characterizing individual brain health and detecting deviations from normative aging trajectories [61, 80]. Prior work has shown that sMRI captures morphological atrophy patterns associated with aging [81, 82], whereas resting-state fMRI provides complementary information on age-related changes in functional network organization [36, 83, 84]. However, most previous studies have either focused on a single modality or adopted simple concatenation-based multimodal fusion [42, 85], which may not fully exploit the complementary nature of functional and structural information.

In this study, we proposed a novel AMAge-Net framework by integrating time series fMRI and sMRI data. The model employs a hierarchical GAT to capture functional features and utilizes a 3D DenseNet121 to extract multi-scale structural information. Structural features provide anatomical context, while functional features encode inter-regional interaction patterns. To achieve effective integration between modalities, we first employ a Multi-Head Cross-Attention mechanism to guide the functional features to attend to local anatomical structures, enabling deep interaction at the local level. This design enables the model to integrate static anatomical constraints with dynamic functional signals, rather than treating the two modalities as independent feature sets. Subsequently, a Gated Fusion mechanism is applied to adaptively combine the global structural representations with the cross-attention features, dynamically modulating the contribution of each modality. This fusion strategy effectively captures and integrates multi-level brain features, providing richer and more discriminative representations for brain age prediction

### Performance Benchmarking and Fusion Strategy Impact

To ensure a fair and representative comparison, the baseline models were selected based on two complementary criteria. Specifically, we included recent graph-based and attention-based models that are representative of current state-of-the-art approaches in brain age prediction and multimodal fusion, such as graph neural networks, GNN [30], GCN [50], BC-GCN [52] and attention- or transformer-based multimodal models MFFormer([58] and CTransformer [59]). In parallel, we compared our method with well-established and widely adopted benchmark architectures with distinctive structural designs, including ResNet [39], EfficientNet [40], and 3D-CNN [41]. These models have consistently demonstrated strong and stable performance across numerous neuroimaging and brain age prediction studies, serving as widely accepted reference points in the field. Together with the selected recent network-based methods, this comparison strategy enables a fair and informative evaluation, avoiding redundant variants of closely related architectures while reflecting both the evolution and methodological diversity of existing modeling approaches.

Our model consistently outperformed all baselines across MAE, RMSE, *R*^2^, and PCC. Notably, models relying exclusively on functional connectivity data performed substantially worse than those using structural features, reaffirming the dominant role of sMRI in brain age prediction. However, not all multimodal models yielded improvements. For instance, MFFormer [58], despite integrating sMRI and fMRI data, underperformed compared to some high-performing models that use only sMRI. This suggests that the choice and design of fusion strategy play a more critical role than simply including multiple data types.

Our model’s significant improvement over 3D-CNN 3D-CNN [41](the strongest model using only structural features) in both prediction accuracy and correlation further supports this idea. By enabling cross-modal interactions through attention and adapting fusion weights via gating, our method captures more relevant and individualized brain aging patterns. The attention-based fusion not only boosts performance but also adds interpretability, allowing insights into modality importance in different prediction contexts [56, 86].

Furthermore, an additional validation was conducted on the OASIS-3 dataset. The consistent performance trends across different models and all variants in the ablation study strengthen the evidence that AMAge-Net captures modality-invariant aging-related features rather than dataset-specific patterns. The results demonstrate that AMAge-Net maintains strong predictive performance on the OASIS-3 dataset, confirming that the proposed model generalizes beyond the Cam-CAN cohort. Notably, Rossel et al. recently evaluated their cosmology-inspired method on both Cam-CAN and OASIS-3 [87], achieving an MAE of 5.9 and 3.1, respectively. While their OASIS-3 result is lower, it was based on 869 sessions from only 378 subjects, which may partially reflect the benefits of multiple scans per subject. In contrast, AMAge-Net achieved comparable performance (MAE = 5.09 on Cam-CAN, 4.29 on OASIS-3) using strictly non-overlapping subjects, providing an alternative evaluation that emphasizes subject-level generalization. Furthermore, large-scale work by Moguilner et al. [88] has demonstrated that brain age predictions are sensitive to population-level differences such as geographic and socioeconomic factors. Together, these findings emphasize the importance of cross-cohort validation, and our dual-dataset evaluation highlights the capacity of AMAge-Net to generalize across diverse populations and imaging protocols.

### Model-Derived Salient Regions and Neurobiological Significance

Beyond predictive performance, a critical advantage of our framework is its ability to provide interpretability through modality-specific saliency analysis.

For the fMRI branch, attention-weight-derived importance scores were used to highlight importantly contributed brain regions, including the superior medial frontal gyrus, the middle temporal pole, the inferior temporal gyrus, and the posterior cingulate cortex. These regions indicated the functional changing patterns of aging. The superior medial prefrontal cortex plays a critical role in higher-order cognitive processes, particularly in planning, decision-making, and executive control. Functional alterations in this region have been associated with age-related declines in goal-directed behavior and the ability to flexibly adapt to changing environmental demands, underscoring its importance in the context of cognitive aging [89]. The middle temporal pole showed a significant contribution to brain age prediction, aligning with its established role in semantic memory, emotional processing, and social cognition [90–96]. Besides, the association analysis between the functional connectivity and age also validated the significant correlation involving the inferior temporal gyrus and the postcentral gyrus. The contribution of these brain regions to the brain age prediction may reflect neurodegenerative changes, underscoring their potential as a neuroimaging marker for age-related cognitive and affective decline.

For the sMRI branch, gradient-based saliency maps emphasized the hippocampus, cuneus, paracentral lobule, and postcentral. Compared to the brain regions emphasized by the fMRI branch, these highly contributed regions in the sMRI branch reveal structural changes of the brain with aging. The hippocampus has been consistently identified across numerous studies as a critical structure for memory formation, consolidation, and retrieval, as well as for broader cognitive functions [97]. Age-related structural atrophy and functional decline in the hippocampus have been linked to impairments in episodic memory, spatial navigation, and learning capacity, and are considered early indicators of pathological aging such as mild cognitive impairment and Alzheimer’s disease [98, 99]. These findings underscore its central role in maintaining cognitive health and highlight the hippocampus as a key target for studying mechanisms of brain aging and potential interventions to preserve cognitive function. Besides, structural MRI studies have demonstrated that age-related cortical thinning in the precentral and postcentral gyri, traditionally associated with motor and somatosensory processing, is also coupled with declines in cognitive domains such as processing speed and executive function [100, 101].

In addition, several regions—including the posterior cingulate cortex, precuneus, and thalamus were salient across both modalities, suggesting convergent structural–functional contributions to brain age estimation. These findings are well aligned with existing literature on the neurobiology of aging [36, 62, 102, 103]. The posterior cingulate and precuneus are key hubs of the default mode network and are among the earliest and most consistently reported regions showing reduced connectivity and cortical thinning with age [104, 105]. The thalamus is known to undergo volumetric decline and decreased network centrality in aging [64, 106], with implications for slowed processing speed and cognitive decline. Similarly, the insula and frontal association cortices have been shown to exhibit age-dependent grey matter loss [65, 66, 83] and altered functional coupling [107, 108], potentially reflecting compensatory recruitment during cognitive tasks.

In the gender-difference analysis, most top contributing regions from the sMRI branch overlapped with those identified in the overall model without gender stratification, and were highly similar across sexes—most notably the posterior cingulate cortex, precuneus, hippocampus, and thalamus. By contrast, the fMRI branch showed greater sex-dependent variability. Overlapping frontal regions identified were the inferior orbitofrontal gyrus, inferior opercular frontal gyrus, and olfactory cortex. These regions are involved in emotion regulation, language production, and olfaction [109–112]. This pattern suggests that sex differences in brain aging may partly reflect variability in socio-emotional and sensory-cognitive networks [113]. Additionally, the motor cortex appeared among the top contributors only in males, whereas the fusiform and occipital regions were prominent only in females. This pattern implies that sensorimotor network changes may disproportionately influence male brain-aging trajectories, while alterations in visual and face-processing systems may play a larger role in female-specific cognitive aging [113, 114].

From a modeling perspective, the convergence of salient regions across modalities strengthens confidence in the biological validity of our predictions. Rather than relying solely on statistical accuracy, our framework identifies neuroanatomically plausible patterns consistent with well-established aging mechanisms [115, 116]. This dual emphasis on predictive accuracy and interpretability is crucial for translational applications of brain age modeling—particularly in detecting accelerated aging in at-risk populations [117, 118], monitoring disease progression [117, 119, 120], or assessing the impact of lifestyle interventions on brain health [121].

### Limitations and Future Work

Some limitations of the present study should be acknowledged. First, the current model focuses on resting-state fMRI and structural MRI. Incorporating additional modalities, such as diffusion MRI or task-based fMRI, as well as non-imaging information (e.g., cognitive or genetic data), may further enhance brain age estimation and provide a more comprehensive characterization of individual aging trajectories. Besides, while the proposed attention-guided fusion strategy improves interpretability by highlighting salient brain regions, the saliency measures remain model-derived and do not establish causal relationships between brain regions and aging processes. Future studies could integrate longitudinal data, causal modeling approaches, or task-based imaging to better distinguish between compensatory and degenerative mechanisms underlying brain aging.

Furthermore, the current study primarily focuses on brain age estimation in healthy individuals and has not yet explored the application of the proposed framework to patients with neurodegenerative disorders such as cognitive impairment and Alzheimer’s disease. Brain age deviation is considered a potential biomarker for disease progression, particularly reflecting accelerated brain aging in AD [122, 123] and mild cognitive impairment [96]. Future work could extend the model to clinical datasets, incorporating pathological status for training and validation, to evaluate the model’s sensitivity and ability to detect pathological brain aging. This would help explore its potential clinical value for early diagnosis and prognosis assessment.

## Supporting information

supplementray for the age prediction

## Supporting information

**S1 Appendix. Details in methods and implementation** The details of participants and data preprocessing, implementation, evaluation metrics used in this study, and construction of the whole-brain functional network.

**S2 Appendix. Participants and Data Preprocessing of OASIS-3** The data protocal of OASIS-3 and their preprocessing.

**S1. Figure. Performance on OASIS-3**. The predicted and chronological age of AMAge-Net on the OASIS-3 dataset.

**S2. Figure. Ablation study across different models on OASIS-3**. The results of the ablation study across different models. The performance metrics include MAE, RMSE, *R*^2^, and PCC.

**S1. Table. Brain atlas**. The anatomical regions defined in each hemisphere and their label in the automated anatomical labelling atlas

**S2. Table. Method comparsion on OASIS-3**. Comparison of our proposed method with other methods in brain age estimation.

**S3. Table. fMRI**. Contribution of brain regions from fMRI.

**S4. Table. sMRI**. Contribution of brain regions from sMRI.

**S5. Table. The correlation between functional connectivity and age**. The top 200 functional connectivity links ranked by the strength of correlation with chronological age from the whole-brain correlation analysis

**S6. Table. Gender difference from fMRI**. The top 23 brain regions with the highest contributions in the fMRI data for male and female subjects in the brain age prediction.

**S7. Table. Gender difference from sMRI**. The top 23 brain regions with the highest contributions in the sMRI data for male and female subjects in the brain age prediction.

**S8. Table. Ablation study on OASIS-3**. Quantitative results of the ablation studies compare different models and feature combinations on OASIS-3.

## Data and Code Availability Statement

The neuroimaging and behavioural data used in this study were obtained from the Cambridge Centre for Ageing and Neuroscience (CamCAN) repository (https://camcan-archive.mrc-cbu.cam.ac.uk) and Open Access Series of Imaging Studies 3 (OASIS-3), which provides open access to multimodal neuroimaging and cognitive data across the adult lifespan. Researchers can apply for access through the CamCAN and OASIS-3 data portal. The code used for data preprocessing and analysis in this study is available at Github.

## Acknowledgments

This work was supported by the Australian Research Council (ARC), Future Fellowship (FT200100942), the Juan De La Cierva (JDC2024-055992-I) and Ramón y Cajal Fellowship (RYC2022-035106-I) from FSE/Agencia Estatal de Investigación (AEI), Spanish Ministry of Science and Innovation, the María de Maeztu Program for Units of Excellence in R&D, grant CEX2021-001164-M and Natural Science Foundation of Jiangsu Province, China (Grant No. BK20230422). The funder played no role in study design, data collection, analysis and interpretation of data, or the writing of this manuscript.

## Author Contributions

Z. W.: Conceptualization, Data Curation, Formal Analysis, Methodology, Investigation, Writing – Original Draft Preparation, Writing – Review & Editing. WX. F.: Data Curation, Methodology, Software, Formal Analysis, Writing – Original Draft Preparation. J. H.: Original Draft Preparation, Writing – Review & Editing. L. L. G..: Supervision, Original Draft Preparation, Writing – Review & Editing. KC. W.: Data Curation, Formal Analysis, Methodology, Supervision, Original Draft Preparation, Writing – Review & Editing.

