## supplementray for the age prediction for "Attention-Guided Multimodal Neuroimaging Fusion Network for Modeling Brain Aging Pattern"

### Supplementary Material

Zhuo Wan[1,*], Wanxiang Fu[2], Javed Hossain[1], Leonardo L. Gollo[3,4], Kaichao Wu[3,4,*]

1. School of Artificial Intelligence, Nanjing University of Information Science and Technology, Nanjing 210044, China
2. School of Computer Science, Nanjing University of Information Science and Technology, Nanjing 210044, China
3. Monash University, Melbourne, VIC 3800, Australia
4. Institute for Cross-Disciplinary Physics and Complex Systems, IFISC (UIB–CSIC), University of the Balearic Islands, Palma de Mallorca, 07122, Spain

* Corresponding author

**Methods in detail**

**Participants and Data Preprocessing Cam-CAN**

The Cambridge Centre for Ageing and Neuroscience (Cam-CAN) (1,2) Stage 2 cohort study is a large-scale (approx. N = 700), multi-modal (MRI, MEG, and behavioural), cross-sectional, population-based adult lifespan (18–87 years old) investigation of the neural underpinnings of successful cognitive ageing. The project is an interdisciplinary collaboration involving researchers with expertise in cognitive psychology, cognitive neuroscience, psychiatry, engineering, and public health. A key focus of the Cam-CAN project is integrative analysis across domains of cognition and measures of neural structure, function, and connectivity, with the goal of understanding how neurocognitive systems adapt in order to overcome age-related changes.

The Stage 2 repository contains MRI, MEG, and behavioural data from 656 participants aged 18–87 years old. All data are labelled with unique project IDs. Data were quality-control checked by semi-automated scripts monitored by the Cam-CAN methods team. All analysis scripts (including aa modules and recipes) are stored in the repository and can therefore be viewed by any user. Matlab scripts are also available to query the repository and compile data cross-referenced by participant identifiers. Using semi-automated Matlab and Linux shell scripts, raw data were pulled from various sources (testing laptops, MRI and MEG data servers) into a central location. Once there, further automated scripts identified new raw data and submitted them to the appropriate processing scripts. Behavioural data were analysed by custom Matlab scripts; MRI and MEG data were processed using Automatic Analysis (aa) (3) pipelines and modules which called relevant functions from neuroimaging analysis software and toolboxes (SPM12, Wellcome Department of Imaging Neuroscience, London, UK; FSL(4); Freesurfer, Martinos Center for Biomedical Imaging, Massachusetts General Hospital, Boston, MA, USA; in-house code).

All MRI datasets were collected at a single site (MRC-CBSU) using a 3 T Siemens TIM Trio scanner with a 32-channel head coil. Participants were scanned in a single 1-hour session. Before scanning, physiological measurements were taken, and two behavioural experiments were run. In the scanner, memory foam cushions were used for comfort and to minimise head movement. Although the collected MRI data included both structural (T1, T2, DWI, and MTI) and functional modalities (resting-state, movie-watching, and sensorimotor tasks), we focused on T1-weighted structural MRI and resting-state functional MRI (rs-fMRI) data. A high resolution 3D T1-weighted structural image is acquired using a Magnetization Prepared RApid Gradient Echo (MPRAGE) sequence with the following parameters: Repetition Time (TR) =2250 milleseconds; Echo Time (TE) =2.99 milliseconds; Inversion Time (TI) =900 milliseconds; flip angle =9 degrees; field of view (FOV) =256mm x 240mm x 192mm; voxel size =1mm isotropic; GRAPPA acceleration factor =2; acquisition time of 4 minutes and 32 seconds. To assess intrinsic (passive) aspects of neural connectivity, T2*-weighted fMRI data are acquired while participants rest with their eyes shut using a Gradient-Echo Echo-Planar Imaging (EPI) sequence. A total of 261 volumes are acquired, each containing 32 axial slices (acquired in descending order), slice thickness of 3.7 mm with an interslice gap of 20% (for whole brain coverage including cerebellum; TR =1970 milliseconds; TE =30 milliseconds; flip angle =78 degrees; FOV =192 mm × 192 mm; voxel-size =3 mm × 3 mm × 4.44 mm) and acquisition time of 8 minutes and 40 seconds.

**Participants and Data Preprocessing of OASIS**

As a complement, this work further evaluated the proposed method using neuroimaging data from the Open Access Series of Imaging Studies, Phase 3 (OASIS-3). Specifically, OASIS-3 is a large, publicly available neuroimaging dataset designed to support studies of normal aging and Alzheimer's disease. The Imaging data of OASIS-3 were acquired across multiple sessions using standardized protocols and underwent rigorous quality control, making OASIS-3 well-suited for assessing the generalizability of the proposed brain age prediction models across datasets with different population characteristics and acquisition settings. We evaluated our developed method with T1 structural and resting-state functional images from 533 normal subjects with ages ranging from 42 to 95 years.

Detailed information about the dataset and imaging protocols is available on the OASIS website (https://sites.wustl.edu/oasisbrains/home/oasis-3/).

OASIS-3 MRI data were collected through the Knight Alzheimer's Disease Research Imaging Program at Washington University in St. Louis, MO, USA (LaMontagne et al., 2019), with a Siemens TIM Trio 3T scanner (Siemens Medical Solutions USA, Inc.). All participants were scanned in the supine position, with head motion minimized using foam padding. In some scanning sessions, a vitamin E capsule was placed over the left temple to provide an external marker for anatomical lateralization. A 16-channel head coil was used for all acquisitions.

The high-resolution T1-weighted structural images were acquired using a three-dimensional magnetization-prepared rapid gradient-echo (3D MPRAGE) sequence with the following parameters: inversion time (TI) = 1000 ms, repetition time (TR) = 2400 ms, echo time (TE) = 3.08 ms, flip angle = 8°, and isotropic voxel size of 1 $mm^3$. Resting-state functional images of OASIS-3 were acquired as part of the OASIS-3 protocol on Siemens 3 T scanners with a TR of 2200 ms, TE of 27 ms, $4 \times 4 \times 4$ $mm^3$ isotropic voxels, a $64 \times 64$ imaging matrix, ~36 slices, and approximately 174 volumes per run (~6 min).

resting-state fMRI data were preprocessed using FSL, including (1) removal of the first four volumes, (2) slice timing correction, (3) motion correction with motion parameter estimation, and (4) mean functional image computation. The functional images were then linearly registered to the corresponding structural images, followed by linear and non-linear registration of the structural images to the MNI standard space. The resulting deformation fields were applied to the functional images, which were then subjected to temporal band-pass filtering (0.01 – 0.1 Hz) and spatial smoothing (full-width at half-maximum, FWHM = 6 mm). After preprocessing, fMRI data from each subject were parcellated using the AAL (Automated Anatomical Labeling) atlas. The atlas was resampled to match the functional image space, and standardized regional time series were extracted using NiftiLabelsMasker. Functional connectivity matrices were then constructed by computing Pearson correlation coefficients between regional time series.

Structural T1-weighted images were obtained from the official FreeSurfer preprocessing pipeline provided by OASIS-3, which includes bias field correction, skull stripping, tissue segmentation, cortical surface reconstruction, and anatomically guided brain parcellation. The resulting functional and structural representations were used for subsequent multimodal fusion modeling.

**Metrics used in this study**

a) MAE (Mean absolute error)：The MAE is used to measure the average magnitude of the absolute differences between the predicted and true age values. It provides a straightforward interpretation of prediction accuracy in the same units as age. The calculation formula is as follows:

$$MAE = \frac{1}{n}\sum_{i=1}^{n} |y_i - \hat{y}_i|$$

Where $y_i$ and $\hat{y}_i$ denote the true and predicted age for subject i, respectively, and n is the total number of subjects.

b) $R^2$ (R-squared)：The $R^2$ quantifies the proportion of variance in the true age values that is predictable from the predicted values. An $R^2$ value closer to 1 indicates better model performance, while negative values suggest the model performs worse than simply predicting the mean age. The calculation formula is as follows:

$$R^2 = 1 - \frac{\sum_{i=1}^{n}(y_i - \hat{y}_i)^2}{\sum_{i=1}^{n}(y_i - \bar{y})^2}$$

where $\bar{y}$ is the mean of the true ages.

c) RMSE (Root Mean Squared Error): The RMSE evaluates the square root of the average squared differences between predicted and actual ages. It penalizes larger errors more heavily than MAE and shares the same unit as the age values. The calculation formula is as follows:

$$RMSE = \sqrt{\frac{1}{n}\sum_{i=1}^{n}(y_i - \hat{y}_i)^2}$$

d) PCC (Pearson Correlation Coefficient) : The PCC measures the linear correlation between the predicted and true ages. A value close to 1 indicates a strong positive linear relationship. The calculation formula is as follows:

$$PCC = \frac{\sum_{i=1}^{n}(y_i - \bar{y})(\hat{y}_i - \bar{\hat{y}})}{\sqrt{\sum_{i=1}^{n}(y_i - \bar{y})^2}\sqrt{\sum_{i=1}^{n}(\hat{y}_i - \bar{\hat{y}})^2}}$$

Where $\bar{y}$ and $\bar{\hat{y}}$ denote the means of the true and predicted ages, respectively.

### Implementation setup

The proposed model was implemented using PyTorch and trained on an NVIDIA GeForce RTX 4090 GPU. Model optimization was performed using the Adam optimizer with an initial learning rate of $1\times10^{-4}$.. To enhance convergence efficiency and training stability, a performance-based learning rate scheduler was employed. The learning rate was automatically reduced by a predefined factor when the validation performance plateaued for a set number of epochs, until reaching a minimum threshold of $1\times10^{-}6$. A five-fold cross-validation strategy was employed to ensure robustness and generalizability. All subjects were randomly divided into five subsets of equal size. In each fold, four subsets were used for training and the remaining one for testing.

The model was trained for 50 epochs with a batch size of 8. Training one fold of AMAge-Net required approximately 2 hours, and the total number of trainable parameters is 11.53 million. After model warm-up, the average inference time was approximately 8 ms per subject, indicating that the proposed framework enables efficient prediction and is computationally feasible for large-scale applications.

### Construction of the Whole-Brain Functional Network

After preprocessing, the resting-state functional magnetic resonance imaging (fMRI) data were parcellated into 116 regions of interest (ROIs) using the AAL atlas (5). Based on previous literature on brain age prediction, this study focused on the cerebrum and excluded the cerebellum from the analysis. The mean voxel signals within each cerebral region were computed to obtain the corresponding time series. For each participant, Pearson cross-correlations of the blood oxygen level - dependent signals were calculated between all pairs of cerebral regions, resulting in a whole-brain functional connectivity network (90 $\times$ 90 regions with 4,050 edges which are functional connectivity links).

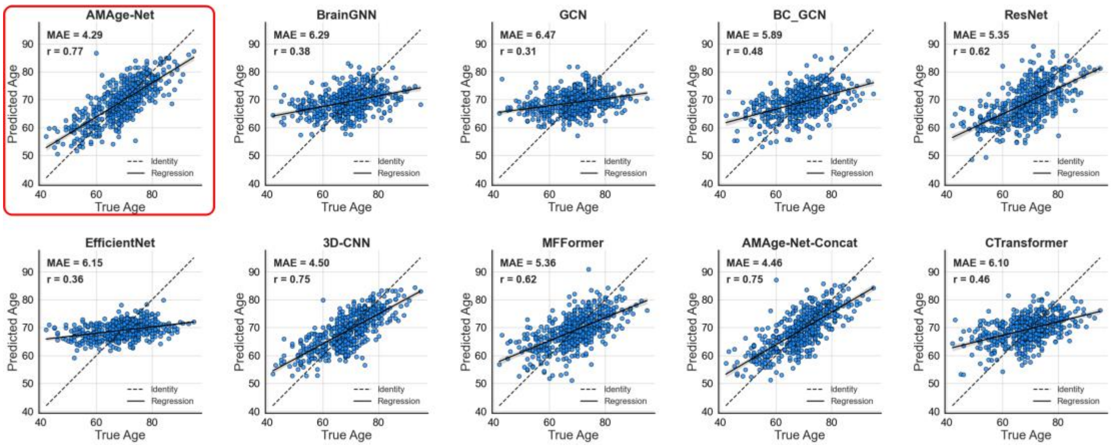


**S1. Figure.** The predicted and chronological age of AMAge-Net on the OASIS-3 dataset. In comparison with unimodal and alternative multimodal methods (e.g., BrainGNN, GCN, MFFormer, and concatenation-based fusion), AMAge-Net consistently achieves a lower MAE (4.29) and higher correlation (r = 0.77), indicating superior generalization performance on an independent cohort.

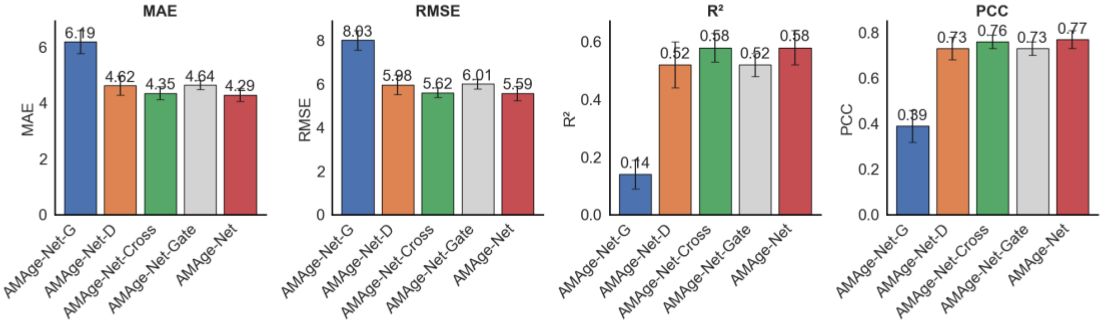


**S2. Figure**. The results of the ablation study across different models. The performance metrics include MAE, RMSE, $R^2$, and PCC.

**Table S1.** The anatomical regions defined in each hemisphere and their label in the automated anatomical labelling atlas. Column 4 provides a set of possible abbreviations for the anatomical descriptions.

| **NO.** | **Anatomical description** | **Label** | **Abbreviation** |
|---|---|---|---|
| 1,2 | Precental gyrus | Precentral | PreCG |
| 3,4 | Superior frontal gyrus, dorsolateral | Frontal_Sup | SFGdor |
| 5,6 | Superior frontal gyrus, orbital part | Frontal_Sup_Orb | ORBsup |
| 7,8 | Middle frontal gyrus | Frontal_Mid | MFG |
| 9,10 | Middle frontal gyrus, orbital part | Frontal_Mid_Orb | ORBmid |

| | | | |
|---|---|---|---|
| 11,12 | Inferior frontal gyrus, opercular part | Frontal_Inf_Oper | IFGoperc |
| 13,14 | Inferior frontal gyrus, triangular part | Frontal_Inf_Tri | IFGtriang |
| 15,16 | Inferior frontal gyrus, orbital part | Frontal_Inf_Orb | ORBinf |
| 17,18 | Rolandic operculum | Rolandic_Oper | ROL |
| 19,20 | Supplementary motor area | Supp_Motor_Area | SMA |
| 21,22 | Olfactory cortex | Olfactory | OLF |
| 23,24 | Superior frontal gyrus, medial | Frontal_Sup_Medial | SFGmed |
| 25,26 | Superior frontal gyrus, medial orbital | Frontal_Med_Orb | ORBsupmed |
| 27,28 | Gyrus rectus | Rectus | REC |
| 29,30 | Insula | Insula | INS |
| 31,32 | Anterior cingulate and paracingulate gyri | Cingulum_Ant | ACG |
| 33,34 | Median cingulate and paracingulate gyri | Cingulum_Mid | DCG |
| 35,36 | Posterior cingulate gyrus | Cingulum_Post | PCG |
| 37,38 | Hippocampus | Hippocampus | HIP |
| 39,40 | Parahippocampal gyrus | ParaHippocampal | PHG |
| 41,42 | Amygdala | Amygdala | AMYG |
| 43,44 | Precuneus fissure and surrounding cortex | Calcarine | CAL |
| 45,46 | Cuneus | Cuneus | CUN |
| 47,48 | Lingual gyrus | Lingual | LING |
| 49,50 | Superior occipital gyrus | Occipital_Sup | SOG |
| 51,52 | Middle occipital gyrus | Occipital_Mid | MOG |
| 53,54 | Inferior occipital gyrus | Occipital_Inf | IOG |
| 55,56 | Fusiform gyrus | Fusiform | FFG |
| 57,58 | Postcentral gyrus | Postcentral | PoCG |
| 59,60 | Superior parietal gyrus | Parietal_Sup | SPG |
| 61,62 | Inferior parietal, but supramarginal and angular gyri | Parietal_Inf | IPL |
| 63,64 | Supramarginal gyrus | SupraMarginal | SMG |
| 65,66 | Angular gyrus | Angular | ANG |
| 67,68 | Precuneus | Precuneus | PCUN |
| 69,70 | Paracentral lobule | Paracentral_Lobule | PCL |
| 71,72 | Caudate nucleus | Caudate | CAU |
| 73,74 | Lenticular nucleus, putamen | Putamen | PUT |
| 75,76 | Lenticular nucleus, pallidum | Pallidum | PAL |
| 77,78 | Thalamus | Thalamus | THA |
| 79,80 | Heschl gyrus | Heschl | HES |
| 81,82 | Superior temporal gyrus | Temporal_Sup | STG |
| 83,84 | Temporal pole: superior temporal gyrus | Temporal_Pole_Sup | TPOsup |
| 85,86 | Middle temporal gyrus | Temporal_Mid | MTG |
| 87,88 | Temporal pole: middle temporal gyrus | Temporal_Pole_Mid | TPOmid |
| 89,90 | Inferior temporal gyrus | Temporal_Inf | ITG |

**Table S2.** Comparison of our proposed method with other methods in brain age estimation on OASIS-3.

| Methods | Modality | MAE | RMSE | R2 | PCC |
|---|---|---|---|---|---|
| BC_GCN | fMRI | 5.89 | 7.65 | 0.22 | 0.48 |
| GCN | fMRI | 6.47 | 8.33 | 0.08 | 0.31 |
| BrainGNN | fMRI | 6.29 | 8.11 | 0.13 | 0.38 |
| Resnet | fMRI | 5.35 | 6.97 | 0.35 | 0.62 |
| EfficientNet | fMRI | 6.15 | 8.11 | 0.13 | 0.36 |
| 3D CNN | sMRI | 4.50 | 5.72 | 0.56 | 0.75 |
| MFFormer | sMRI+fMRI | 5.36 | 6.86 | 0.37 | 0.62 |
| CTransformer | sMRI+fMRI | 6.10 | 7.75 | 0.20 | 0.46 |
| AMAge-Net-Concat | sMRI+fMRI | 4.46 | 5.78 | 0.55 | 0.75 |
| **AMAge-Net (Ours)** | sMRI+fMRI | **4.29** | **5.59** | **0.58** | **0.77** |

**Table S3.** The **top 200** functional connectivity links ranked by the strength of correlation with chronological age from the whole-brain correlation analysis. Only connections with significant correlations (p<0.05 FDR corrected) are shown.

| Region 1 | Region 2 | r value | p value | Region 1 | Region 2 | r value | p value |
|---|---|---|---|---|---|---|---|
| Rolandic_Oper_R | Temporal_Sup_L | -0.469 | 2.12E-33 | Occipital_Inf_L | Postcentral_R | -0.317 | 3.98E-15 |
| Rolandic_Oper_L | Temporal_Sup_L | -0.428 | 2.67E-27 | Cingulum_Mid_R | Temporal_Pole_Sup_L | -0.317 | 3.98E-15 |
| Rolandic_Oper_R | Caudate_L | -0.429 | 2.67E-27 | Precentral_R | Temporal_Inf_L | -0.316 | 4.84E-15 |
| Rolandic_Oper_R | Caudate_R | -0.405 | 4.34E-24 | Temporal_Pole_Sup_L | Temporal_Mid_R | -0.315 | 6.13E-15 |
| Insula_L | Temporal_Pole_Sup_L | -0.397 | 3.90E-23 | Heschl_L | Temporal_Pole_Sup_R | -0.315 | 6.66E-15 |
| Frontal_Med_Orb_L | Cingulum_Post_L | -0.395 | 5.79E-23 | Rolandic_Oper_R | Insula_L | -0.315 | 6.74E-15 |
| Rolandic_Oper_R | Temporal_Pole_Sup_R | -0.393 | 8.74E-23 | Precentral_R | Paracentral_Lobule_R | -0.314 | 7.54E-15 |
| Cuneus_R | Lingual_R | -0.391 | 1.57E-22 | Postcentral_R | Parietal_Inf_R | -0.314 | 7.54E-15 |
| Rolandic_Oper_L | Temporal_Pole_Sup_R | -0.389 | 2.12E-22 | Frontal_Sup_Medial_R | Temporal_Pole_Sup_L | -0.314 | 8.57E-15 |
| Occipital_Mid_L | Fusiform_L | -0.387 | 4.16E-22 | Rolandic_Oper_R | Occipital_Inf_R | -0.313 | 1.05E-14 |
| Rolandic_Oper_L | Caudate_L | -0.387 | 4.16E-22 | Frontal_Inf_Oper_R | Temporal_Pole_Sup_R | -0.313 | 1.10E-14 |
| Rolandic_Oper_L | Temporal_Pole_Sup_L | -0.384 | 7.82E-22 | Frontal_Sup_Orb_R | Frontal_Inf_Tri_R | -0.313 | 1.10E-14 |
| Occipital_Sup_L | Fusiform_L | -0.382 | 1.46E-21 | Postcentral_L | Temporal_Inf_R | -0.312 | 1.11E-14 |
| Rolandic_Oper_R | Heschl_R | -0.377 | 5.06E-21 | Precentral_L | Temporal_Inf_L | -0.312 | 1.16E-14 |
| Rolandic_Oper_R | Temporal_Pole_Sup_L | -0.376 | 6.51E-21 | Frontal_Sup_Orb_L | Cingulum_Post_L | -0.312 | 1.19E-14 |
| Cingulum_Ant_R | Temporal_Pole_Sup_L | -0.373 | 1.48E-20 | Rectus_R | Cingulum_Post_L | -0.312 | 1.27E-14 |
| Rolandic_Oper_L | Caudate_R | -0.373 | 1.49E-20 | Postcentral_R | Temporal_Sup_L | -0.312 | 1.31E-14 |
| Insula_R | Temporal_Pole_Sup_L | -0.372 | 1.78E-20 | Frontal_Inf_Tri_R | Frontal_Inf_Orb_R | -0.311 | 1.40E-14 |
| Cingulum_Ant_L | Temporal_Pole_Sup_L | -0.371 | 2.31E-20 | Precuneus_L | Temporal_Inf_L | -0.309 | 2.09E-14 |
| Rolandic_Oper_R | Temporal_Sup_R | -0.369 | 3.35E-20 | Frontal_Inf_Tri_R | Temporal_Pole_Sup_R | -0.309 | 2.16E-14 |
| Cuneus_R | Lingual_L | -0.369 | 3.4E-20 | Frontal_Mid_R | Frontal_Mid_Orb_R | -0.309 | 2.23E-14 |
| Cuneus_L | Lingual_R | -0.368 | 3.95E-20 | Cuneus_R | Occipital_Inf_R | -0.309 | 2.37E-14 |
| Rolandic_Oper_R | Heschl_L | -0.368 | 4.03E-20 | Cingulum_Mid_L | Temporal_Pole_Sup_L | -0.308 | 2.71E-14 |
| Frontal_Inf_Tri_R | Temporal_Pole_Sup_L | -0.366 | 6.68E-20 | Occipital_Mid_R | Occipital_Inf_R | -0.307 | 3.31E-14 |
| Lingual_R | Occipital_Sup_R | -0.365 | 9.84E-20 | Cingulum_Ant_R | Temporal_Pole_Sup_R | -0.307 | 3.41E-14 |
| Cuneus_L | Lingual_L | -0.362 | 1.81E-19 | Rolandic_Oper_L | Insula_R | -0.307 | 3.67E-14 |
| Occipital_Mid_L | Temporal_Inf_R | -0.360 | 3.11E-19 | Rolandic_Oper_R | Cingulum_Ant_R | -0.306 | 3.82E-14 |

| | | | | | | | |
|---|---|---|---|---|---|---|---|
| Occipital_Mid_R | Fusiform_R | -0.360 | 3.29E-19 | Frontal_Sup_Orb_L | Precuneus_L | -0.306 | 3.86E-14 |
| Frontal_Inf_Tri_L | Temporal_Pole_Sup_L | -0.359 | 3.53E-19 | Rolandic_Oper_R | Temporal_Mid_R | -0.306 | 4.40E-14 |
| Frontal_Inf_Oper_L | Temporal_Pole_Sup_L | -0.359 | 3.79E-19 | Rolandic_Oper_L | Temporal_Inf_L | -0.306 | 4.40E-14 |
| Occipital_Mid_R | Fusiform_L | -0.358 | 4.88E-19 | Supp_Motor_Area_L | Temporal_Pole_Sup_L | -0.305 | 4.91E-14 |
| Occipital_Sup_L | Fusiform_R | -0.357 | 5.61E-19 | Rolandic_Oper_R | Cingulum_Mid_L | -0.305 | 4.91E-14 |
| Occipital_Sup_R | Occipital_Inf_R | -0.356 | 6.9E-19 | Fusiform_L | Postcentral_L | -0.305 | 5.53E-14 |
| Parietal_Sup_L | Temporal_Inf_R | -0.356 | 8.29E-19 | Frontal_Mid_R | Temporal_Pole_Sup_L | -0.304 | 5.83E-14 |
| Rolandic_Oper_L | Fusiform_L | -0.355 | 8.59E-19 | Calcarine_L | Temporal_Pole_Sup_L | -0.304 | 5.87E-14 |
| Precentral_R | Occipital_Inf_R | -0.355 | 9.06E-19 | Frontal_Sup_Orb_R | Cingulum_Post_R | -0.304 | 5.93E-14 |
| Occipital_Sup_R | Fusiform_L | -0.355 | 1.04E-18 | Rolandic_Oper_R | Pallidum_R | -0.304 | 6.07E-14 |
| Occipital_Sup_R | Fusiform_R | -0.354 | 1.19E-18 | Cuneus_L | Fusiform_R | -0.304 | 6.20E-14 |
| Occipital_Mid_L | Fusiform_R | -0.353 | 1.41E-18 | Frontal_Inf_Tri_L | Temporal_Inf_R | -0.304 | 6.20E-14 |
| Occipital_Mid_L | Temporal_Inf_L | -0.353 | 1.44E-18 | Occipital_Sup_L | Temporal_Inf_L | -0.304 | 6.33E-14 |
| Frontal_Inf_Oper_L | Temporal_Inf_L | -0.353 | 1.58E-18 | Frontal_Sup_Orb_L | Cingulum_Post_R | -0.304 | 6.33E-14 |
| Parietal_Inf_L | Temporal_Inf_L | -0.352 | 1.73E-18 | Cingulum_Mid_L | Temporal_Inf_L | -0.303 | 6.99E-14 |
| Frontal_Inf_Tri_L | Temporal_Inf_L | -0.350 | 3.27E-18 | Cingulum_Ant_L | Cingulum_Mid_R | -0.303 | 7.43E-14 |
| Rectus_L | Cingulum_Post_L | -0.348 | 4.99E-18 | Rolandic_Oper_R | Fusiform_L | -0.302 | 8.52E-14 |
| Occipital_Inf_R | Postcentral_R | -0.348 | 5.35E-18 | Cingulum_Mid_R | Temporal_Inf_R | -0.302 | 8.87E-14 |
| Precentral_R | Temporal_Mid_L | -0.346 | 8.18E-18 | Cuneus_L | Temporal_Pole_Sup_L | -0.302 | 8.90E-14 |
| Frontal_Med_Orb_R | Cingulum_Post_L | -0.346 | 8.63E-18 | Occipital_Inf_L | Temporal_Inf_R | -0.302 | 9.83E-14 |
| Frontal_Inf_Oper_R | Temporal_Pole_Sup_L | -0.341 | 2.44E-17 | Fusiform_L | Postcentral_R | -0.301 | 1.02E-13 |
| Parietal_Sup_L | Temporal_Inf_L | -0.341 | 2.64E-17 | Rolandic_Oper_R | Pallidum_L | -0.301 | 1.04E-13 |
| Occipital_Sup_R | Occipital_Inf_L | -0.340 | 3.25E-17 | Precentral_L | Temporal_Inf_R | -0.301 | 1.19E-13 |
| Insula_L | Temporal_Pole_Sup_R | -0.340 | 3.37E-17 | Rolandic_Oper_L | Insula_L | -0.300 | 1.40E-13 |
| Precentral_R | Temporal_Sup_L | -0.340 | 3.37E-17 | Frontal_Sup_L | Temporal_Pole_Sup_L | -0.299 | 1.49E-13 |
| Occipital_Mid_R | Temporal_Inf_R | -0.339 | 3.86E-17 | Rolandic_Oper_R | Temporal_Inf_R | -0.299 | 1.54E-13 |
| Rolandic_Oper_R | Insula_R | -0.339 | 4.45E-17 | Frontal_Sup_Orb_L | Parietal_Inf_L | -0.299 | 1.57E-13 |
| Parietal_Sup_R | Temporal_Inf_R | -0.338 | 5.01E-17 | Occipital_Mid_R | Temporal_Inf_L | -0.299 | 1.73E-13 |
| Occipital_Sup_R | Temporal_Inf_R | -0.338 | 5.21E-17 | Occipital_Mid_L | Occipital_Inf_L | -0.299 | 1.76E-13 |
| Lingual_L | Occipital_Sup_R | -0.338 | 5.42E-17 | Rolandic_Oper_L | Cingulum_Mid_R | -0.298 | 1.80E-13 |
| Lingual_R | Occipital_Sup_L | -0.337 | 6.13E-17 | Precentral_R | Fusiform_R | -0.298 | 1.84E-13 |
| Frontal_Sup_Medial_L | Temporal_Pole_Sup_L | -0.337 | 6.64E-17 | Frontal_Mid_R | Temporal_Inf_L | -0.298 | 1.92E-13 |
| Rolandic_Oper_R | Cingulum_Ant_L | -0.336 | 6.95E-17 | Occipital_Mid_R | Occipital_Inf_L | -0.298 | 1.99E-13 |
| Occipital_Sup_L | Occipital_Inf_L | -0.335 | 1.06E-16 | Precuneus_L | Temporal_Inf_R | -0.298 | 2.10E-13 |
| Precentral_R | Fusiform_L | -0.333 | 1.70E-16 | Rolandic_Oper_L | Cingulum_Ant_R | -0.297 | 2.31E-13 |
| Rolandic_Oper_L | Paracentral_Lobule_R | -0.332 | 1.92E-16 | Frontal_Mid_Orb_R | Parietal_Inf_R | -0.297 | 2.36E-13 |
| Occipital_Sup_L | Temporal_Inf_R | -0.332 | 1.92E-16 | Rolandic_Oper_R | Temporal_Mid_L | -0.297 | 2.55E-13 |
| Insula_R | Temporal_Pole_Sup_R | -0.331 | 2.57E-16 | Cingulum_Mid_L | Lingual_R | -0.296 | 2.62E-13 |
| Rolandic_Oper_L | Fusiform_R | -0.330 | 2.77E-16 | Cingulum_Mid_L | Temporal_Inf_R | -0.296 | 2.76E-13 |
| Heschl_L | Temporal_Pole_Sup_L | -0.330 | 2.77E-16 | Cingulum_Mid_R | Lingual_R | -0.296 | 2.80E-13 |
| Parietal_Inf_L | Temporal_Inf_R | -0.330 | 3.06E-16 | Frontal_Sup_R | Temporal_Pole_Sup_L | -0.296 | 3.05E-13 |
| Cingulum_Ant_L | Cingulum_Mid_L | -0.329 | 3.40E-16 | Frontal_Med_Orb_L | Cingulum_Post_R | -0.295 | 3.23E-13 |
| Rolandic_Oper_L | Occipital_Inf_R | -0.329 | 3.45E-16 | Cingulum_Mid_R | Temporal_Inf_L | -0.295 | 3.42E-13 |
| Precentral_R | Occipital_Inf_L | -0.329 | 3.50E-16 | Temporal_Pole_Sup_L | Temporal_Mid_L | -0.295 | 3.58E-13 |
| Rolandic_Oper_R | Cingulum_Mid_R | -0.329 | 3.61E-16 | Frontal_Sup_Orb_L | Parietal_Sup_L | -0.295 | 3.58E-13 |
| Cuneus_L | Fusiform_L | -0.328 | 4.41E-16 | Frontal_Mid_Orb_L | Parietal_Inf_L | -0.295 | 3.63E-13 |
| Frontal_Inf_Oper_R | Temporal_Inf_L | -0.327 | 5.03E-16 | Rolandic_Oper_L | Temporal_Sup_R | -0.295 | 3.64E-13 |
| Frontal_Inf_Oper_R | Temporal_Inf_R | -0.327 | 5.95E-16 | Rolandic_Oper_R | Fusiform_R | -0.294 | 3.78E-13 |
| Precentral_R | Temporal_Inf_R | -0.327 | 5.95E-16 | Cingulum_Ant_R | Cingulum_Mid_R | -0.294 | 3.92E-13 |
| Precentral_L | Temporal_Mid_L | -0.325 | 8.63E-16 | Frontal_Sup_Orb_R | Cingulum_Post_L | -0.294 | 3.94E-13 |
| Lingual_L | Occipital_Sup_L | -0.325 | 8.91E-16 | Precentral_L | Fusiform_L | -0.294 | 4.54E-13 |
| Rolandic_Oper_L | Postcentral_R | -0.325 | 9.26E-16 | Lingual_R | Temporal_Pole_Sup_L | -0.293 | 5.00E-13 |
| Occipital_Sup_L | Occipital_Inf_R | -0.324 | 9.86E-16 | Rolandic_Oper_R | Postcentral_R | -0.293 | 5.42E-13 |
| Rolandic_Oper_L | Temporal_Inf_R | -0.324 | 1.07E-15 | Postcentral_L | Temporal_Inf_L | -0.293 | 5.42E-13 |

| | | | | | | | |
|---|---|---|---|---|---|---|---|
| Postcentral_R | Temporal_Inf_R | -0.323 | 1.25E-15 | Cingulum_Mid_R | Temporal_Pole_Sup_R | -0.292 | 6.32E-13 |
| Frontal_Sup_Orb_R | Frontal_Mid_R | -0.323 | 1.26E-15 | Frontal_Med_Orb_R | Cingulum_Post_R | -0.291 | 6.93E-13 |
| Precentral_R | Temporal_Pole_Sup_L | -0.322 | 1.79E-15 | Precentral_L | Occipital_Inf_R | -0.291 | 6.99E-13 |
| Precentral_R | Lingual_L | -0.322 | 1.80E-15 | Lingual_R | Temporal_Inf_R | -0.291 | 6.99E-13 |
| Occipital_Mid_L | Temporal_Pole_Sup_L | -0.321 | 1.84E-15 | Rolandic_Oper_L | Hippocampus_R | -0.291 | 7.09E-13 |
| Frontal_Mid_L | Temporal_Pole_Sup_L | -0.321 | 1.87E-15 | Rolandic_Oper_L | Supp_Motor_Area_L | -0.291 | 7.09E-13 |
| Precentral_R | Lingual_R | -0.321 | 1.89E-15 | Fusiform_L | Parietal_Sup_L | -0.291 | 7.18E-13 |
| Frontal_Mid_L | Temporal_Inf_L | -0.321 | 2.16E-15 | Frontal_Sup_Orb_R | Parietal_Inf_L | -0.291 | 7.21E-13 |
| Occipital_Inf_R | Postcentral_L | -0.320 | 2.31E-15 | Rolandic_Oper_L | Heschl_R | -0.291 | 7.45E-13 |
| Frontal_Inf_Oper_L | Temporal_Inf_R | -0.320 | 2.51E-15 | Frontal_Inf_Oper_L | Temporal_Pole_Sup_R | -0.291 | 7.91E-13 |
| Cuneus_R | Fusiform_R | -0.319 | 2.97E-15 | Precentral_L | Temporal_Pole_Sup_L | -0.290 | 9.17E-13 |
| Cuneus_R | Fusiform_L | -0.319 | 2.97E-15 | Angular_L | Temporal_Inf_L | -0.289 | 9.75E-13 |
| Postcentral_R | Temporal_Mid_L | -0.319 | 2.97E-15 | Frontal_Sup_L | Temporal_Inf_L | -0.289 | 9.90E-13 |
| Frontal_Inf_Tri_R | Temporal_Inf_L | -0.319 | 3.20E-15 | Insula_R | Cingulum_Mid_R | -0.289 | 9.99E-13 |
| Precentral_R | Temporal_Mid_R | -0.319 | 3.20E-15 | Temporal_Sup_R | Temporal_Pole_Sup_L | -0.289 | 1.01E-12 |
| Calcarine_R | Temporal_Pole_Sup_L | -0.318 | 3.43E-15 | Rolandic_Oper_L | Postcentral_L | -0.288 | 1.18E-12 |
| Occipital_Mid_L | Occipital_Inf_R | -0.318 | 3.43E-15 | Cingulum_Post_L | Temporal_Inf_L | -0.288 | 1.28E-12 |
| Paracentral_Lobule_L | Paracentral_Lobule_R | -0.318 | 3.64E-15 | Frontal_Sup_Orb_R | Frontal_Mid_L | -0.288 | 1.30E-12 |
| Frontal_Inf_Tri_R | Temporal_Inf_R | -0.318 | 3.88E-15 | Lingual_R | Postcentral_R | -0.287 | 1.47E-12 |

**Table S4.** This table presents the relative contributions of brain regions from **fMRI** data to the prediction results in the brain age prediction task. The importance values were computed using attention weights, reflecting the influence of each region in the prediction process. Brain region names follow the standard nomenclature of the AAL atlas.

| **Brain_Region** | **Importance** | **Brain_Region** | **Importance** |
|---|---|---|---|
| Thalamus_R | 1 | Frontal_Mid_Orb_L | 0.562 |
| Frontal_Sup_Medial_L | 0.993 | Temporal_Pole_Mid_L | 0.556 |
| Insula_L | 0.945 | SupraMarginal_R | 0.549 |
| Temporal_Pole_Mid_R | 0.922 | Frontal_Mid_R | 0.549 |
| Caudate_L | 0.882 | Cingulum_Mid_L | 0.536 |
| Frontal_Inf_Orb_R | 0.824 | Precuneus_R | 0.529 |
| Rectus_L | 0.824 | Postcentral_L | 0.523 |
| Frontal_Mid_Orb_R | 0.810 | Insula_R | 0.516 |
| Paracentral_Lobule_R | 0.810 | Temporal_Sup_R | 0.516 |
| Cingulum_Post_R | 0.804 | Frontal_Inf_Tri_R | 0.510 |
| Frontal_Inf_Oper_R | 0.784 | Cuneus_R | 0.503 |
| Temporal_Inf_R | 0.778 | Fusiform_L | 0.490 |
| Precentral_R | 0.771 | Frontal_Sup_Orb_R | 0.477 |
| Rectus_R | 0.758 | Cuneus_L | 0.471 |
| Occipital_Mid_R | 0.745 | Cingulum_Ant_L | 0.464 |
| Rolandic_Oper_R | 0.745 | Pallidum_R | 0.458 |
| SupraMarginal_L | 0.745 | Lingual_R | 0.458 |
| Frontal_Med_Orb_R | 0.745 | Lingual_L | 0.451 |
| Temporal_Pole_Sup_L | 0.745 | Temporal_Mid_R | 0.451 |

| | | | |
|---|---|---|---|
| Thalamus_L | 0.732 | Temporal_Sup_L | 0.418 |
| Occipital_Sup_L | 0.732 | Frontal_Mid_L | 0.405 |
| Frontal_Med_Orb_L | 0.725 | Putamen_L | 0.399 |
| Postcentral_R | 0.725 | ParaHippocampal_R | 0.399 |
| ParaHippocampal_L | 0.712 | Hippocampus_R | 0.379 |
| Supp_Motor_Area_R | 0.699 | Calcarine_R | 0.372 |
| Cingulum_Mid_R | 0.693 | Paracentral_Lobule_L | 0.366 |
| Cingulum_Post_L | 0.693 | Rolandic_Oper_L | 0.353 |
| Calcarine_L | 0.686 | Precentral_L | 0.340 |
| Precuneus_L | 0.673 | Parietal_Sup_L | 0.333 |
| Pallidum_L | 0.673 | Frontal_Inf_Tri_L | 0.333 |
| Cingulum_Ant_R | 0.667 | Caudate_R | 0.327 |
| Angular_R | 0.654 | Occipital_Inf_R | 0.320 |
| Parietal_Inf_R | 0.634 | Frontal_Sup_Medial_R | 0.314 |
| Occipital_Inf_L | 0.627 | Temporal_Pole_Sup_R | 0.307 |
| Heschl_L | 0.614 | Putamen_R | 0.301 |
| Olfactory_L | 0.608 | Fusiform_R | 0.275 |
| Frontal_Sup_Orb_L | 0.608 | Amygdala_R | 0.261 |
| Temporal_Mid_L | 0.595 | Occipital_Mid_L | 0.242 |
| Frontal_Inf_Oper_L | 0.595 | Parietal_Inf_L | 0.229 |
| Frontal_Sup_L | 0.589 | Frontal_Sup_R | 0.176 |
| Hippocampus_L | 0.582 | Amygdala_L | 0.137 |
| Parietal_Sup_R | 0.575 | Temporal_Inf_L | 0.092 |
| Supp_Motor_Area_L | 0.575 | Heschl_R | 0.092 |
| Occipital_Sup_R | 0.569 | Frontal_Inf_Orb_L | 0.078 |
| Olfactory_R | 0.652 | Angular_L | 0 |

**Table S5.** This table presents the relative contributions of brain regions from **sMRI** data to the prediction results in the brain age prediction task. The importance values were derived from saliency maps computed on sMRI inputs, reflecting the influence of each region in the prediction process. Higher values indicate greater impact. Brain region names follow the standard nomenclature of the AAL atlas.

| **Brain_Region** | **Importance** | **Brain_Region** | **Importance** |
|---|---|---|---|
| Cingulum_Post_R | 1 | Angular_L | 0.280 |
| Precuneus_R | 0.831 | Fusiform_L | 0.278 |
| Calcarine_R | 0.729 | Hippocampus_L | 0.276 |
| Cingulum_Post_L | 0.725 | ParaHippocampal_L | 0.271 |
| Cuneus_R | 0.717 | Caudate_L | 0.256 |
| Paracentral_Lobule_R | 0.684 | Frontal_Mid_R | 0.254 |
| Cuneus_L | 0.684 | Temporal_Mid_R | 0.252 |
| Thalamus_R | 0.675 | Postcentral_L | 0.251 |

| | | | |
|---|---|---|---|
| Lingual_R | 0.663 | Parietal_Inf_L | 0.245 |
| Calcarine_L | 0.662 | Temporal_Mid_L | 0.224 |
| Precuneus_L | 0.661 | Precentral_L | 0.212 |
| Occipital_Sup_L | 0.655 | SupraMarginal_L | 0.206 |
| Parietal_Sup_R | 0.627 | Frontal_Sup_L | 0.205 |
| Heschl_R | 0.623 | Frontal_Sup_Medial_L | 0.201 |
| Occipital_Mid_L | 0.611 | Rectus_R | 0.201 |
| Paracentral_Lobule_L | 0.610 | Temporal_Inf_L | 0.197 |
| Postcentral_R | 0.592 | Occipital_Mid_R | 0.193 |
| Hippocampus_R | 0.588 | Temporal_Pole_Sup_R | 0.192 |
| Lingual_L | 0.570 | Olfactory_L | 0.189 |
| Parietal_Inf_R | 0.531 | Heschl_L | 0.186 |
| Parietal_Sup_L | 0.520 | Temporal_Sup_L | 0.185 |
| Precentral_R | 0.515 | Frontal_Inf_Tri_L | 0.183 |
| ParaHippocampal_R | 0.507 | Frontal_Sup_Medial_R | 0.181 |
| Insula_R | 0.480 | Frontal_Inf_Tri_R | 0.179 |
| Rolandic_Oper_R | 0.474 | Frontal_Mid_L | 0.178 |
| Supp_Motor_Area_R | 0.470 | Temporal_Inf_R | 0.175 |
| SupraMarginal_R | 0.464 | Frontal_Inf_Oper_L | 0.168 |
| Cingulum_Mid_R | 0.464 | Rectus_L | 0.167 |
| Occipital_Sup_R | 0.464 | Temporal_Pole_Sup_L | 0.164 |
| Fusiform_R | 0.462 | Frontal_Inf_Orb_L | 0.161 |
| Occipital_Inf_L | 0.460 | Temporal_Pole_Mid_L | 0.160 |
| Caudate_R | 0.436 | Rolandic_Oper_L | 0.159 |
| Thalamus_L | 0.426 | Putamen_L | 0.159 |
| Putamen_R | 0.425 | Insula_L | 0.157 |
| Angular_R | 0.420 | Frontal_Inf_Orb_R | 0.139 |
| Temporal_Sup_R | 0.419 | Amygdala_L | 0.128 |
| Amygdala_R | 0.380 | Pallidum_L | 0.125 |
| Cingulum_Ant_R | 0.368 | Frontal_Med_Orb_R | 0.112 |
| Supp_Motor_Area_L | 0.365 | Frontal_Med_Orb_L | 0.096 |
| Cingulum_Mid_L | 0.351 | Occipital_Inf_R | 0.093 |
| Frontal_Inf_Oper_R | 0.342 | Frontal_Sup_Orb_R | 0.091 |
| Cingulum_Ant_L | 0.323 | Frontal_Sup_Orb_L | 0.085 |
| Frontal_Sup_R | 0.312 | Temporal_Pole_Mid_R | 0.077 |
| Pallidum_R | 0.311 | Frontal_Mid_Orb_L | 0.052 |
| Olfactory_R | 0.286 | Frontal_Mid_Orb_R | 0 |

**Table S6.** This table presents the top 24 brain regions with the highest contributions in the **fMRI** data for male and female subjects in the brain age prediction experiment. The left and right columns display the results for **males** and **females**, respectively, allowing for a direct comparison of sex-specific differences in regional contributions.

| Rank | Brain Region(M) | Importance(M) | Brain Region(F) | Importance(F) |
|---|---|---|---|---|
| 1 | Frontal_Mid_Orb_L | 1 | Fusiform_L | 1 |
| 2 | Supp_Motor_Area_L | 0.984 | Frontal_Inf_Oper_R | 0.991 |
| 3 | Hippocampus_L | 0.976 | Temporal_Pole_Mid_R | 0.955 |
| 4 | ParaHippocampal_L | 0.951 | Frontal_Inf_Tri_L | 0.910 |
| 5 | Lingual_L | 0.935 | Parietal_Inf_R | 0.883 |
| 6 | Rolandic_Oper_L | 0.911 | Hippocampus_R | 0.856 |
| 7 | Lingual_R | 0.886 | Rectus_R | 0.856 |
| 8 | Temporal_Inf_L | 0.870 | Temporal_Mid_L | 0.847 |
| 9 | Frontal_Sup_L | 0.821 | Occipital_Inf_R | 0.829 |
| 10 | Supp_Motor_Area_R | 0.821 | Precentral_R | 0.820 |
| 11 | Rolandic_Oper_R | 0.813 | Olfactory_L | 0.811 |
| 12 | Precentral_L | 0.813 | Paracentral_Lobule_L | 0.802 |
| 13 | Frontal_Inf_Oper_L | 0.797 | Parietal_Inf_L | 0.784 |
| 14 | Amygdala_R | 0.772 | Thalamus_L | 0.775 |
| 15 | Frontal_Inf_Orb_L | 0.772 | Pallidum_L | 0.775 |
| 16 | Frontal_Inf_Orb_R | 0.772 | Caudate_R | 0.757 |
| 17 | Temporal_Pole_Mid_R | 0.764 | Frontal_Sup_Orb_R | 0.748 |
| 18 | Frontal_Inf_Tri_R | 0.756 | Calcarine_R | 0.721 |
| 19 | Olfactory_R | 0.756 | Angular_L | 0.712 |
| 20 | Frontal_Inf_Oper_R | 0.748 | Fusiform_R | 0.703 |
| 21 | Olfactory_L | 0.707 | Frontal_Inf_Orb_L | 0.694 |
| 22 | Cingulum_Post_R | 0.700 | Frontal_Sup_Orb_L | 0.685 |
| 23 | Insula_R | 0.691 | Olfactory_R | 0.676 |
| 24 | Angular_R | 0.683 | Cingulum_Post_L | 0.667 |

**Table S7.** This table presents the top 23 brain regions with the highest contributions in the **sMRI** data for male and female subjects in the brain age prediction experiment. The left and right columns display the results for **males** and **females**, respectively, allowing for a direct comparison of sex-specific differences in regional contributions.

| Rank | Brain Region(M) | Importance(M) | Brain Region(F) | Importance(F) |
|---|---|---|---|---|
| 1 | Cingulum_Post_R | 1 | Cingulum_Post_R | 1 |
| 2 | Precuneus_R | 0.800 | Calcarine_R | 0.849 |
| 3 | Thalamus_R | 0.763 | Cingulum_Post_L | 0.837 |
| 4 | Cingulum_Post_L | 0.753 | Hippocampus_R | 0.820 |
| 5 | Paracentral_Lobule_L | 0.717 | Precuneus_R | 0.816 |
| 6 | Paracentral_Lobule_R | 0.716 | Lingual_R | 0.805 |
| 7 | Cuneus_R | 0.706 | Cuneus_R | 0.791 |
| 8 | Calcarine_R | 0.702 | Cuneus_L | 0.775 |
| 9 | Precuneus_L | 0.690 | Heschl_R | 0.775 |

| | | | | |
|---|---|---|---|---|
| 10 | Heschl_R | 0.649 | ParaHippocampal_R | 0.775 |
| 11 | Parietal_Sup_R | 0.648 | Thalamus_R | 0.771 |
| 12 | Lingual_R | 0.638 | Calcarine_L | 0.766 |
| 13 | Postcentral_R | 0.634 | Occipital_Mid_L | 0.748 |
| 14 | Cuneus_L | 0.622 | Occipital_Sup_L | 0.743 |
| 15 | Precentral_R | 0.603 | Amygdala_R | 0.726 |
| 16 | Supp_Motor_Area_R | 0.599 | Parietal_Sup_L | 0.722 |
| 17 | Calcarine_L | 0.590 | Precuneus_L | 0.718 |
| 18 | Hippocampus_R | 0.587 | Thalamus_L | 0.711 |
| 19 | Occipital_Sup_L | 0.577 | Lingual_L | 0.703 |
| 20 | Thalamus_L | 0.576 | Insula_R | 0.674 |
| 21 | Insula_R | 0.539 | Fusiform_R | 0.656 |
| 22 | Supp_Motor_Area_L | 0.535 | Putamen_R | 0.616 |
| 23 | Rolandic_Oper_R | 0.535 | Parietal_Sup_R | 0.609 |

**Table S8**. Quantitative results of the ablation studies compare different models and feature combinations. Evaluations include baseline models using only structural or functional features, models using only Cross-fusion or only Gated fusion, as well as our proposed method

| Methods | Modality | MAE | RMSE | R2 | PCC |
|---|---|---|---|---|---|
| AMAge-Net-G | fMRI | 6.19 | 8.03 | 0.14 | 0.39 |
| AMAge-Net-D | sMRI | 4.62 | 5.98 | 0.52 | 0.73 |
| AMAge-Net-Cross | sMRI+fMRI | 4.35 | 5.62 | 0.58 | 0.76 |
| AMAge-Net-Gate | sMRI+fMRI | 4.64 | 6.01 | 0.52 | 0.73 |
| AMAge-Net (Ours) | sMRI+fMRI | **4.29** | **5.59** | **0.58** | **0.77** |
